# Paternal behavior is controlled by preoptic Trpc5 neurons

**DOI:** 10.64898/2026.03.20.713203

**Authors:** Yongxiang Li, Qingzhuo Liu, Fuhui Wang, Kristine M. McDermott, Mengjie Wang, Yue Deng, Yuxue Yang, Yutian Liu, Jingjing Cheng, Meixin Sun, Xinming Liu, Jinjing Jian, Jiamin Qiu, Xi Wu, Lamei Xue, Tong Zhou, Yongjie Yang, Hailan Liu, Longlong Tu, Benjamin R. Arenkiel, I. Sadaf Farooqi, Yong Xu

## Abstract

Male parenting behavior is highly conserved and activated only after offspring are born. Here we show that the behavioral switch from infanticide in virgin male mice to caregiving in wild-type sires is associated with enhanced expression and activation of Transient receptor potential channel 5 (Trpc5) in estrogen receptor α (Esr1)-expressing neurons in the hypothalamic medial preoptic area (MPOA). While selective deletion of Trpc5 from Esr1^MPOA^ neurons diminishes paternal behavior in sires, overexpression of Trpc5 in these neurons converts the otherwise infanticidal virgin males to exhibit care to pups. Mechanistically, Trpc5-dependent changes in neuronal excitability underlie fatherhood-associated Esr1^MPOA^ neuron plasticity. Notably, Trpc5 overexpression also enhances escape behavior and elicits exploratory diving behavior, suggesting that these neurons control a broader adaptive parenting response in males. These findings establish a Trpc5-dependent MPOA neural signal as a critical regulator of paternal behavior.

## Introduction

In humans, parental care plays a critical role in shaping offspring development, influencing cognitive, emotional, and social outcomes^1–3^. In contrast, early-life abuse or neglect increases the risk of psychiatric disorders and antisocial behavior in children and adolescents ^4,5^. From an evolutionary perspective, parental care is essential for offspring survival across many mammalian species^1,6,7^. While female animals are biologically predisposed to caregiving, male involvement varies greatly. In mice, virgin males typically attack or ignore pups, a behavior thought to enhance reproductive success by keeping females fertile. Moreover, caring for pups demands time and energy and can reduce the virgin male’s overall fitness^8^. Remarkably, after mating and cohabiting with a pregnant female, males undergo a striking behavioral shift, transitioning from infanticide to paternal care.

Recent studies in rodents have provided some insights into the molecular and neural basis of paternal behavior^8–11^. The medial preoptic area (MPOA) of the hypothalamus has emerged as a critical center for the regulation of paternal behavior. Lesions in the MPOA impair paternal responses in California male mice that would naturally provide care to their young^12^. Specific neuronal populations in the MPOA, such as those expressing galanin^13,14^, calcitonin receptor^15^, and bombesin receptor subtype 3^16^ have been shown to influence parental care. Prolactin signaling in this region appears essential for paternal behavior as deletion of the prolactin receptor (Prlr) from the MPOA impairs paternal behaviors in mouse sires^17^. Moreover, inactivating glutamatergic neurons or activating GABAergic neurons in the MPOA reduces pup-directed aggression^18^. Recent work suggests that estrogen receptor α (Esr1)-expressing neurons in the MPOA also regulate multiple social behaviors, including parenting behavior^19^. In particular, inhibition of MPOA Esr1 neurons suppresses pup retrieval behavior in lactating females^19^. In male rats, stimulation of the MPOA with estrogen has been shown to promote paternal behavior^20^. Unlike females, whose hormonal fluctuations across reproductive states are well characterized and known to shape behavioral profiles, males do not exhibit such cyclical endocrine changes, making their capacity for behavioral plasticity particularly intriguing. For instance, while prolactin levels progressively rise from virgin to lactating dams, no prolactin increase is observed in sires^17^, although prolactin receptor signaling in the brain is required for normal paternal care^21^. Therefore, the exact mechanisms that enable male mice to switch from pup-directed aggression to caregiving are still poorly understood.

Transient receptor potential channel 5 (TRPC5) is a brain-expressed non-selective cation channel that regulates neuronal excitability, and has been shown to be activated in response to diverse internal and external sensory cues^22^. We have previously shown that *TRPC5* loss-of-function mutations in humans are associated with food-seeking, obesity, anxiety, autism, and postpartum depression ^23^. These phenotypes were entirely recapitulated in a knock-in mouse model harboring a complete loss-of-function human *Trpc5* mutation (K34del)^23^. Female mothers (dams) carrying the K34del mutation exhibited poor maternal care and depressive-like behavior, which was reversed when Trpc5 expression was restored in the paraventricular nucleus of the hypothalamus (PVH)^23^.

Given its prominent role in mediating instinctive behaviors, we investigated the effects of Trpc5 on paternal behavior, using the knock-in mouse model of human disease (K34del) to identify mechanisms that may be conserved across mammalian species. We found that Trpc5 was expressed by Esr1 neurons in the MPOA and its activation mediated the experience-dependent switch in male pup-directed behavior. We further demonstrated that Trpc5 expression in Esr1^MPOA^ neurons modulates the neurons’ functional properties and mediates fatherhood-associated Esr1^MPOA^ neuron plasticity. Lastly, we found that enhanced Trpc5 expression in Esr1^MPOA^ neurons induces escape behavior and exploratory behavior including pronounced diving in water, suggesting a broader role for Trpc5 in male behavioral responses which may act to defend their offspring.

## Results

### Paternal experience suppresses pup-directed aggression and enhances pup retrieval

Virgin males often commit infanticide to boost reproductive success, but after mating, they temporarily shift to paternal behavior following parturition. To further understand this behavioral transition, we compared pup-directed behaviors between sexually naïve (virgin) C57BL/6J males and experienced sires using a pup-retrieval assay (Fig. 1a). Consistent with previous findings^24^, we found that most virgin males exhibited pup-directed aggression (38.88%), or ignored (55.55%) the pups, reflecting typical infanticidal behavior (Fig. 1b-1c). The virgin male mice were next housed with wild-type (WT) females till three days after parturition, and pup retrieval behavior was assessed. Briefly, dams and their biological pups were removed from the home cage six hours prior to testing, leaving the sires alone. Subsequently, three unfamiliar WT pups (not genetically related to the tested sires) were introduced into the home cage for behavioral assessment. Remarkably, 100% of the sires displayed pup retrieval behavior, with no demonstration of attacking or ignoring pups (Fig. 1d; Extended Data Fig. 1). Compared to virgin males, sires showed a significant reduction in time to retrieve pups, improved retrieval efficiency, and increased number of pups retrieved (Fig. 1e-1g). After the retrieval process, sires spent significantly more time engaging in other paternal behaviors, including crouching over pups, grooming them, building nests, and remaining in the nest, all indicative of increased paternal care (Fig. 1h-1l). These results suggest that fatherhood induces a behavioral switch in C57BL/6J male mice, revealing an experience-dependent plasticity underlying paternal care.

**Figure 1.**
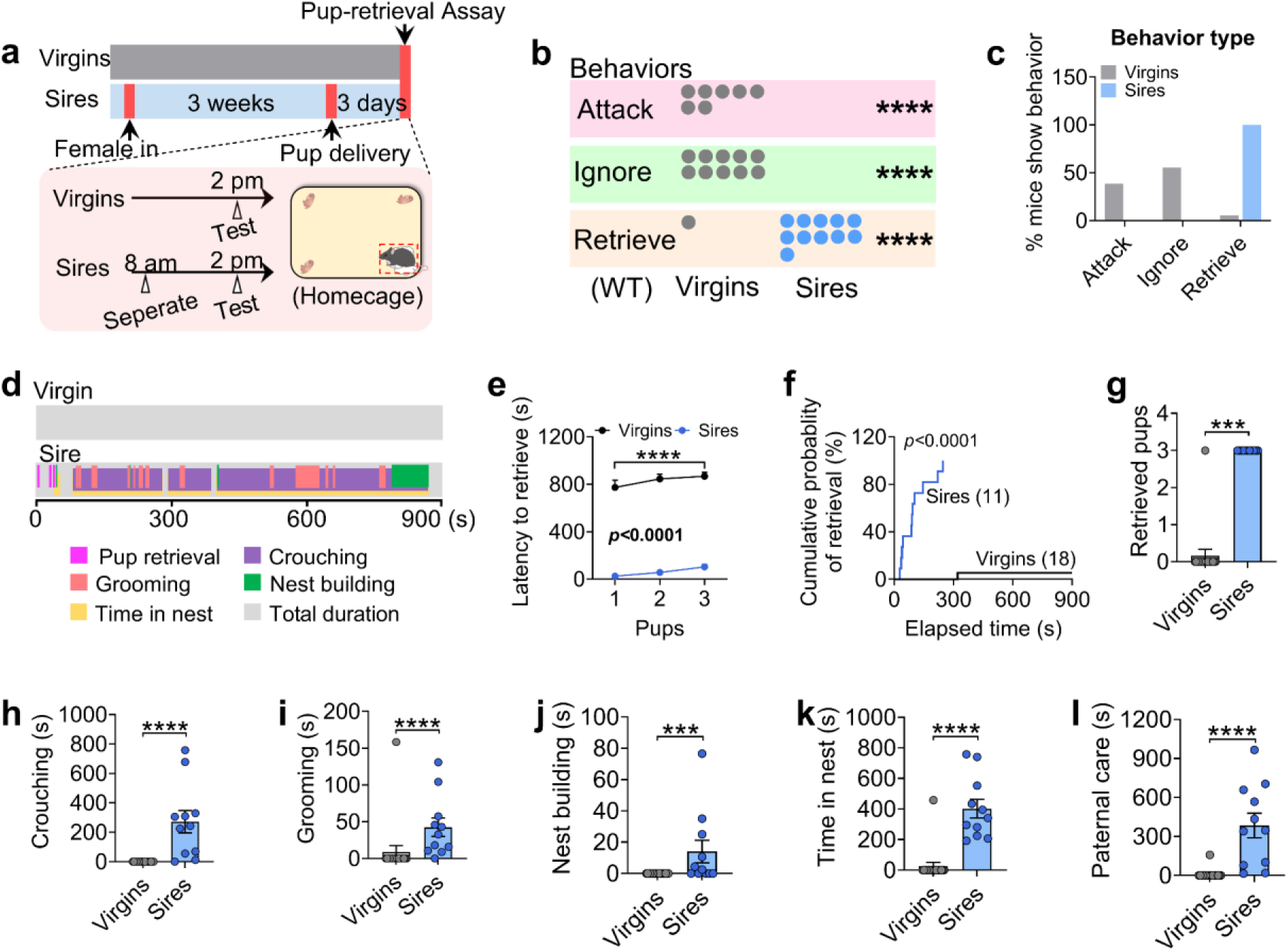
Paternal experience shifts the male behaviors from aggression or neglect to parenting. (a) Illustration showing pup-directed behaviors assessment between WT virgin males and sires co-housed with females following partition (n = 11-18 per group, 12 weeks of age). (b) Dots indicate the number of virgin males and sires that showing attack, ignore or retrieve. (c) Percentage of virgin males and sires showing attack, ignore, or retrieve. (d) Representative behavior raster plots of two individual mouse (virgin male and sire) in the pup-directed behavior. (e) The latency to retrieve pups. (f) Cumulative probability of pup retrieval. (g) The numbers of retrieved pups. (h-l) The duration of crouching pups (h), grooming pups (i), nest building (j), time spent in the nest (k) and paternal care duration (l) (n = 11-18 per group, 12 weeks of age). Data presented as mean ± SEM. p value determined using two-way ANOVA (e), Mann-Whitney test (g,h,i,j,k,l), and chi-squared test (b). ***p < 0.001 and ****p < 0.0001.

### Trpc5 loss-of-function mutations impair paternal behavior in mice

We previously reported that female mice carrying a loss-of-function human mutation in the *Trpc5* gene (K34del; Trpc5^K34del^) exhibit reduced maternal care^23^. Here, we investigated whether the same mutation in male mice affects paternal behavior. To this end, WT and Trpc5^K34del/Y^ virgin males (C57BL/6J background) were mated with WT C57BL/6J female mice (Fig. 2a). Importantly, Trpc5^K34del/Y^ virgin males displayed normal sexual behavior comparable to WT males (Extended Data Fig. 2a-2f). Then, each male was co-housed with the same female mouse throughout pregnancy until three days after parturition. Unlike control males, which reliably retrieved their pups, Trpc5^K34del/Y^ sires largely ignored the pups, and failed to engage in parental caregiving behavior (Fig. 2b-2d; Extended Data Fig. 2g). Notably, Trpc5^K34del/Y^ mutant sires exhibited significantly longer latency to retrieve pups, reduced retrieval efficiency, and decreased number of retrieved pups (Fig. 2e-2g). In addition, Trpc5^K34del/Y^ sires spent significantly less time crouching over pups, grooming, building nests, or remaining in the nest (Fig. 2h-2l). These findings indicate that Trpc5 is required for the expression of paternal behavior in male mice.

**Figure 2.**
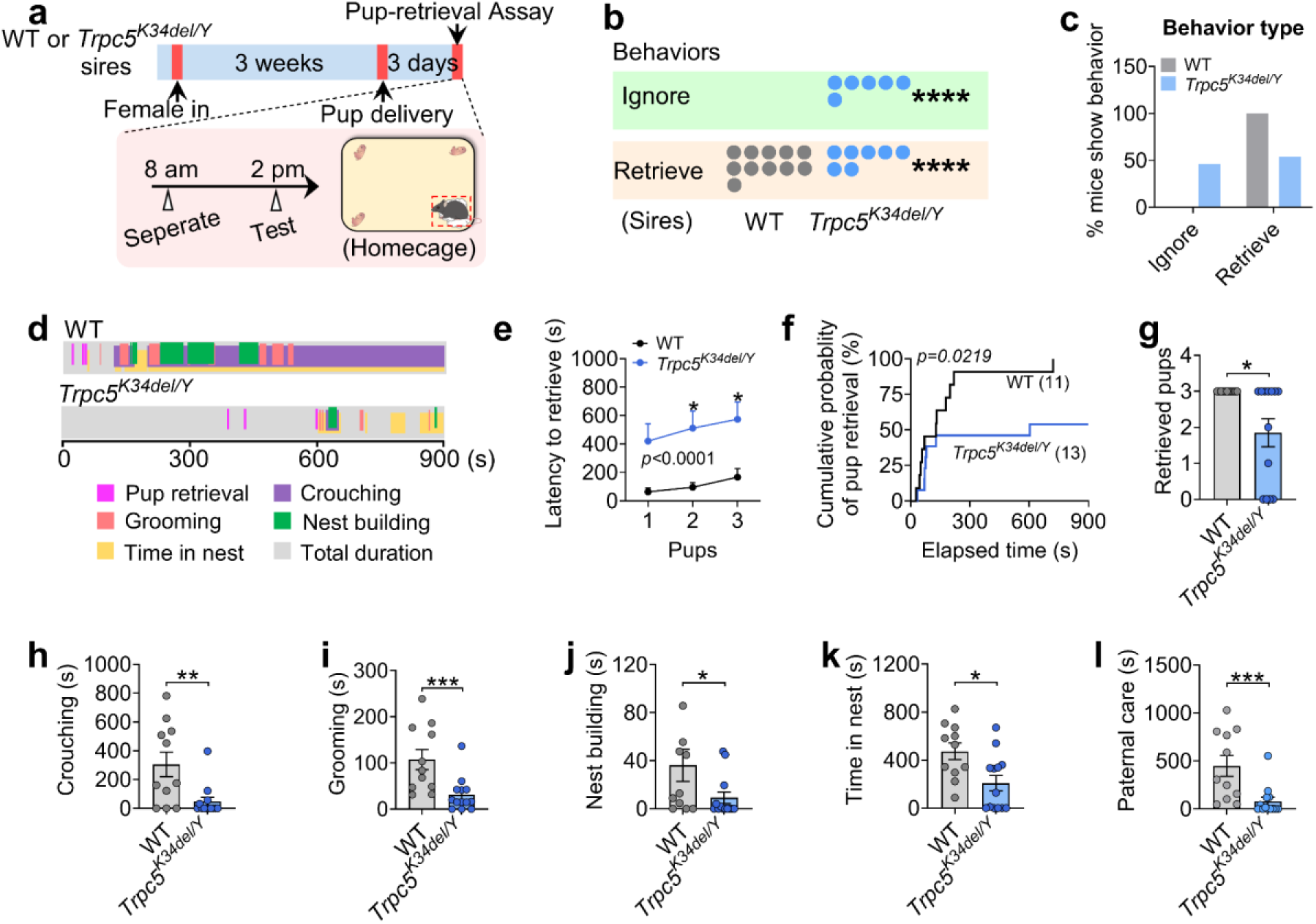
Trpc5^K34del/Y^ sires exhibit impaired paternal behavior. (a) Illustration showing pup-directed behaviors assessment between WT and Trpc5^K34del/Y^ sires co-housed with females following partition (n = 11-13 per group, 16 weeks of age). (b) Dots indicate the number of WT and Trpc5^K34del/Y^ sires that show ignore or retrieve. (c) Percentage of WT and Trpc5^K34del/Y^ sires showing ignore or retrieve. (d) Representative behavior raster plots of two individual mouse (WT and Trpc5^K34del/Y^) in the pup-directed behavior. (e) The latency to retrieve pups. (f) Cumulative probability of pup retrieval. (g) The numbers of retrieved pups. (h-l) The duration of crouching pups (h), grooming pups (i), nest building (j), time spent in the nest (k) and paternal care duration (l) (n = 11-13 per group, 16 weeks of age). Data presented as mean ± SEM. p value determined using two-way ANOVA (e), Mann-Whitney test (g,h,i,j,k,l), and chi-squared test (b). *p < 0.05, **p < 0.01, ***p < 0.001, and ****p < 0.0001.

### Trpc5 expression in Esr1^MPOA^ neurons correlates with paternal behavior in male mice

We next investigated whether Trpc5 expression changes during the transition to fatherhood. Notably, the level of Trpc5 immunoreactivity in the MPOA, a hypothalamic region critical for parental behavior, was significantly elevated in sires that exhibited pup retrieval compared to hostile virgin males (Fig. 3a-3c). Regression analysis revealed that Trpc5+ neuron number and density of Trpc5 immunoreactivity in the MPOA were negatively correlated with pup retrieval latency, and positively correlated with paternal care duration, indicating a strong association between MPOA Trpc5 levels and paternal behavior (Fig. 3d-3g). Trpc5 expression also correlated with key paternal caregiving behaviors, including crouching, grooming, and time spent in the nest (Extended Data Fig. 3a-3f).

**Figure 3.**
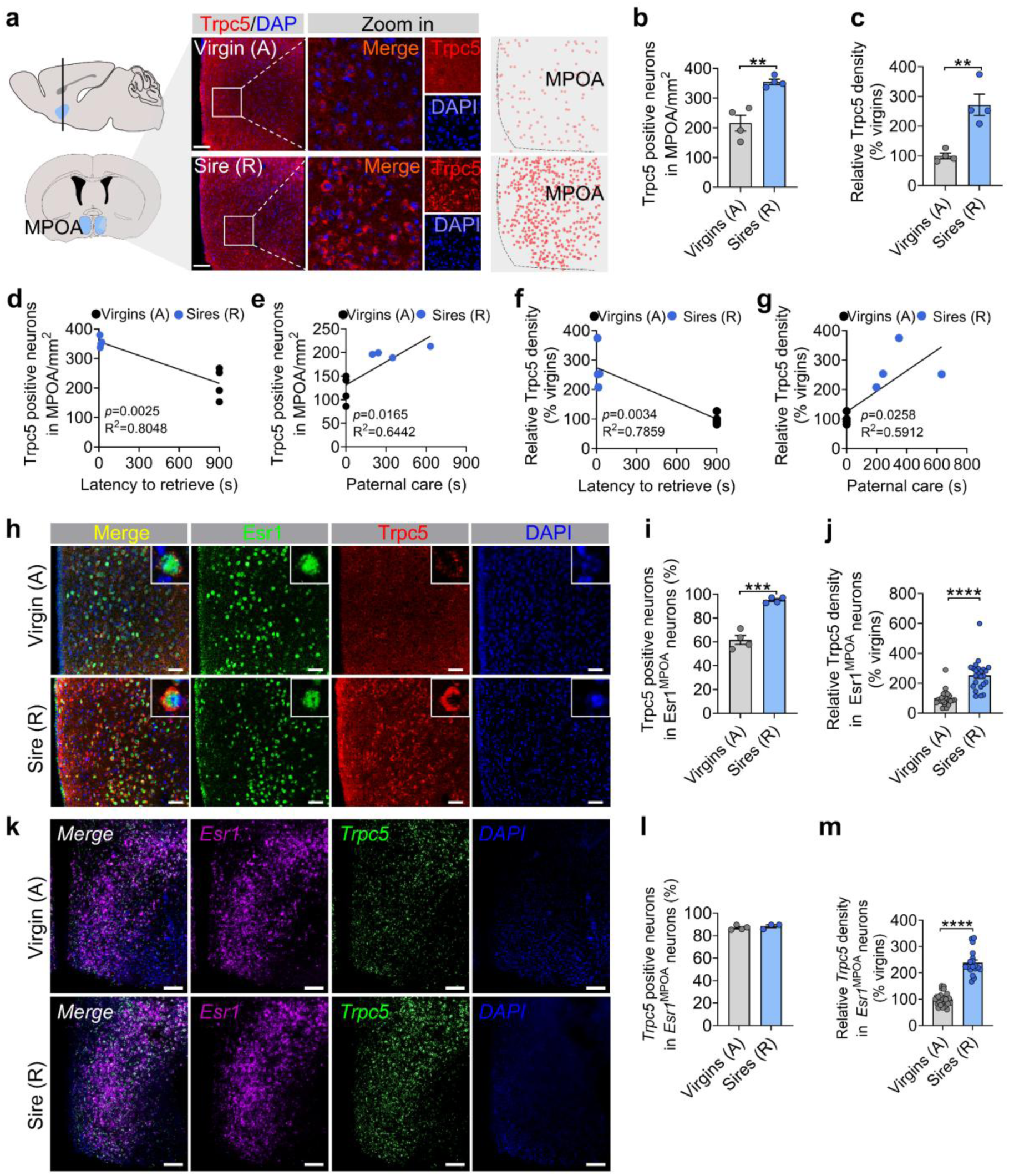
Fatherhood experience increases Trpc5 expression in Esr1 neurons in the MPOA. (a) Immunofluorescence images showing Trpc5 expression in the MPOA of hostile virgin males and pup-retrieving sires (n = 4 per group, 12 weeks of age). Scale bar, 100 μm. (b-c) Quantification of Trpc5+ neuron number (b) and density of Trpc5 immunoreactivity (c) in the MPOA of hostile virgin males and pup-retrieving sires (n = 4 per group, 12 weeks of age). (d-e) Correlation between the number of Trpc5+ neurons in the MPOA and behavioral performance, including latency to retrieve pups (d) and paternal care duration (e). Each dot represents one animal (n = 4 per group, 12 weeks of age). (f-g) Correlation between density of Trpc5 immunoreactivity in the MPOA and latency to retrieve pups (f) and paternal care scores (g). Each dot represents one animal (n = 4 per group, 12 weeks of age). (h-i) Representative images (h) and quantification (i) of the percentage of Esr1^MPOA^ neurons (green) co-expressing Trpc5 (red) in hostile virgin males and pup-retrieving sires (n = 4 mice per group, 12 weeks of age). Scale bar, 100 μm. Nuclei are counterstained with DAPI (blue). (j) Quantification of Trpc5 density within Esr1^MPOA^ neurons (n = 6 neurons per mouse, 4 mice per group, 12 weeks of age). (k-l) Representative RNAscope images (k) and quantification (l) of the percentage of *Esr1*^MPOA^ neurons (red) co-expressing *Trpc5* (green) in hostile virgin males and pup-retrieving sires (n = 3-4 mice per group, 12 weeks of age). Scale bar, 50 μm. Nuclei are counterstained with DAPI (blue). (m) Quantification of *Trpc5* mRNA density within *Esr1*^MPOA^ neurons (n = 6 neurons per mouse, 3-4 mice per group, 12 weeks of age). Data presented as mean ± SEM. p value determined using unpaired t-test (b,c,i,j,l,m). **p < 0.01, ***p < 0.001, and ****p < 0.0001.

Esr1 is broadly expressed in brain regions implicated in social behaviors, and Esr1-expressing neurons in the MPOA have been shown to play a preferential role in regulating parental behavior^19,25^. Given the established role of Esr1^MPOA^ neurons in parental behavior, we further assessed Trpc5 expression within the Esr1^MPOA^ neuron population. Confocal imaging revealed increases in both the number of Trpc5 and Esr1 double-positive neurons in the MPOA and the density of Trpc5 immunoreactivity in Esr1^MPOA^ neurons (Fig. 3h-3j), without changes in total Esr1^MPOA^ neuron number in sires compared to hostile virgin males (Extended Data Fig. 3g). Dual RNAscope experiments confirmed that Trpc5 mRNAs levels in Esr1^MPOA^ neurons were significantly elevated in sires compared to virgin males (Fig. 3k-3m). Lastly, to examine whether exposure to pups activates Esr1^MPOA^ neurons in sires exhibiting retrieval behavior, we assessed c-Fos expression as a marker of neuronal activation following pup re-exposure to their biological sires. Indeed, we observed elevated c-Fos expression in Esr1^MPOA^ neurons following pup re-exposure compared to no re-exposure (Extended Data Fig. 3h). Importantly, we did not detect any difference in Trpc5 expression within the PVH (Extended Data Fig. 4a-4c), the region known to be involved in maternal behavior during these experiments ^23^. Moreover, Trpc5 expression in the PVH was not correlated with pup retrieval latency, crouching, grooming, time spent in the nest, or duration of paternal care (Extended Data Fig. 4d-4m). Together, these findings show that Trpc5 is upregulated in Esr1^MPOA^ neurons and that changes in its expression in these neurons may contribute to the transition to fatherhood.

**Figure 4.**
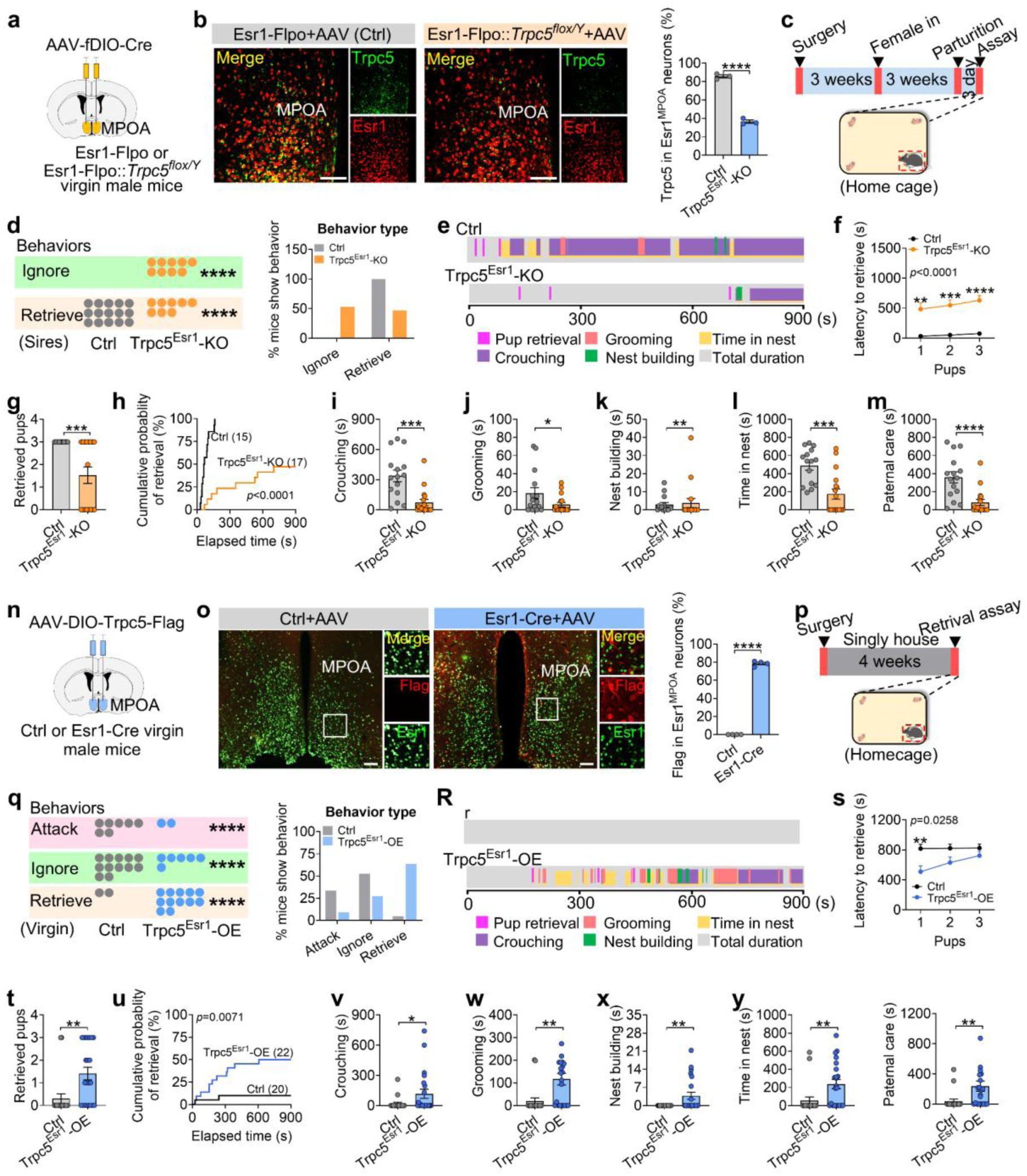
Trpc5 in Esr1^MPOA^ neurons modulates paternal behavior. (a) Schematic diagram of Flpo-dependent AAV vectors used to express Cre recombinase, which were injected into the MPOA of Esr1-Flpo and Esr1-Flpo::Trpc5^flox/Y^ adult male mice. (b) Representative images showing colocalization of Trpc5 (green) and Esr1 (red) neurons in MPOA (left), and quantification of the percentage of Esr1^MPOA^ neurons expressing Trpc5 (right) (n = 3 mice per group, 15 weeks of age). Scale bars, 100 μm. (c) Experimental timeline. (d) Dots indicate the number of controls sires and Trpc5^Esr1^-KO sires that show ignore or retrieve (left) and percentage of control sires and Trpc5^Esr1^-KO sires (right) (n = 15-17 mice per group, 15 weeks of age). (e) Representative behavior raster plots of two individual mouse (control and Trpc5^Esr1^-KO sire) in the pup-directed behavior (n = 15-17 mice per group, 15 weeks of age). (f) The latency to retrieve pups. (g) Cumulative probability of pup retrieval. (h) The numbers of retrieved pups. (i-m) The duration of crouching pups (i), grooming pups (j), nest building (k), time spent in the nest (l) and paternal care duration (m) (n = 15-17 mice per group, 15 weeks of age). (n) Schematic illustration of Cre-dependent AAV vectors used to express AAV-DIO-Trpc5-Flag, which were injected into the MPOA of control and Esr1-Cre adult virgin male mice. (o) Representative images showing colocalization of Esr1 neurons (green) and Flag-tagged Trpc5 expression (red) in the MPOA (left), and quantification of the percentage of Esr1^MPOA^ neurons expressing Flag (right) (n = 4 mice per group, 14 weeks of age). Scale bars, 100 μm. (p) Experimental timeline. (q) Dots indicate the number of control and Trpc5^Esr1^-OE virgin male mice that show attack, ignore or retrieve (left) and percentage of control and Trpc5^Esr1^-OE virgin male mice (right). (n = 20-22 mice per group, 14 weeks of age). (r) Representative behavior raster plots of two individual mouse (control and Trpc5^Esr1^-OE virgin male) in the pup-directed behavior. (s) The latency to retrieve pups. (t) The numbers of retrieved pups. (u) Cumulative probability of pup retrieval. (v-z) The duration of crouching pups (v), grooming pups (w), nest building (x), time spent in the nest (y) and paternal care duration (z) (n = 20-22 mice per group, 14 weeks of age). Data presented as mean ± SEM. p value determined using two-way ANOVA (f,s), unpaired t-test (b), Mann-Whitney test (g,i,j,k,l,m,o,t,v,w,x,y,z), and chi-squared test (d,q). *p < 0.05, **p < 0.01, ***p < 0.001 and ****p < 0.0001.

### Modulation of paternal behavior by Trpc5 expression in Esr1^MPOA^ neurons

Next, we tested whether Trpc5 in Esr1^MPOA^ neurons impacts paternal behavior in male mice. Towards this end, we first deleted Trpc5 selectively from Esr1^MPOA^ neurons in sires and assessed paternal behavior. For this we bilaterally injected AAV-fDIO-Cre into the MPOA of adult Esr1-Flpo (control) and Esr1-Flpo::Trpc5^flox/Y^ (Trpc5^Esr1^-KO) virgin male mice; both groups were mated with WT female mice to produce litters (Fig. 4a). Post hoc immunofluorescence analysis revealed that only 30% of Esr1^MPOA^ neurons expressed Trpc5 in the Trpc5^Esr1^-KO mice, compared to nearly 90% in control sires (Fig. 4b). Although Trpc5^Esr1^-KO mice exhibited normal sexual behavior comparable to control mice (Extended Data Fig. 5a-5f), 52.94% Trpc5^Esr1^-KO sires showed significant deficits in pup retrieval (Fig. 4c-4e; Extended Data Fig. 5g). Specifically, these mice displayed increased retrieval latency, retrieved fewer pups, and had a reduced cumulative probability of retrieval (Fig. 4f-4h). Additionally, other paternal behaviors (crouching, grooming, nest building, and time spent in the nest) were significantly attenuated (Fig. 4i-4m). In contrast, anxiety-like behavior (Extended Data Fig. 6a-6e), depression-like behavior (Extended Data Fig. 6f-6g), and social interactions (Extended Data Fig. 6h-6m) remained unaffected after deletion of Trpc5. However, loss of Trpc5 from Esr1^MPOA^ neurons significantly increased inter-male aggression toward both adult (Extended Data Fig. 6n-6p) and juvenile (Extended Data Fig. 6q-6s) intruders.

Next, we investigated whether pharmacological inhibition of Trpc5 impairs paternal behavior in WT C57BL/6J sires. To this end, we administered the selective Trpc5 inhibitor AC1903 (i.p.; 50 mg kg^-1^), and assessed paternal behaviors. Notably, Trpc5 inhibition led to a significant increase in pup retrieval latency, a decreased cumulative probability of retrieval, and a reduction in the number of pups retrieved (Extended Data Fig. 7a-7e). Other paternal behaviors were also impaired, including crouching, grooming, and time spent in the nest (Extended Data Fig. 7f-7k).

Given that loss of Trpc5 from Esr1^MPOA^ neurons disrupted paternal behavior and increased aggression, we next asked whether enhancing Trpc5 expression in Esr1^MPOA^ neurons would be sufficient to induce paternal behavior and suppress aggression in virgin males. To address this question, we injected AAV-DIO-Trpc5-Flag into the MPOA of Esr1-Cre mice to drive Trpc5 overexpression specifically in Esr1^MPOA^ neurons (Trpc5^Esr1^-OE). WT littermates received the same virus injections and served as controls (Fig. 4n). Immunofluorescence showed that approximately 70% of Esr1^MPOA^ neurons were immunoreactive for Flag in Esr1-Cre mice (Fig. 4o). To further confirm that AAV-DIO-Trpc5-Flag effectively upregulated Trpc5 expression, we performed unilateral injections into the right side of the MPOA in Esr1-Cre virgin males (Extended Data Fig. 8a). This led to a robust increase in Trpc5 expression in Esr1^MPOA^ neurons on the injected side compared to the un-injected side (Extended Data Fig. 8b-8d). These findings confirmed successful overexpression of Trpc5 in Esr1^MPOA^ neurons using a viral approach. Behaviorally, Trpc5 overexpression in virgin male mice induced robust pup retrieval behavior (Fig. 4p-4r; Extended Data Fig. 8e), characterized by reduced retrieval latency, increased numbers of pups retrieved, and cumulative retrieval probability (Fig. 4s-4u). Also, Trpc5^Esr1^-OE mice showed enhanced crouching, grooming, nest building, time spent in the nest, and paternal care duration (Fig. 4v-4z). We did not observe any changes in anxiety-like behaviors in open field tests (Extended Data Fig. 9a-9e). However, Trpc5^Esr1^-OE mice showed a significant reduction in immobility time in the forced swim test (Extended Data Fig. 9f), but not in the tail suspension test (Extended Data Fig. 9g). However, social interaction times were comparable between control and Trpc5^Esr1^-OE mice (Extended Data Fig. 9h-9m). Notably, Trpc5^Esr1^-OE mice also exhibited significantly reduced inter-male aggression toward both adult (Extended Data Fig. 9n-9p) and juvenile intruders (Extended Data Fig. 9q-9s). Collectively, these findings highlight a critical and specific role for Trpc5 in Esr1^MPOA^ neurons in regulating paternal behavior and male-directed aggression.

### Paternal experience remodels the activity of Esr1^MPOA^ neurons via Trpc5

Next, we sought to investigate the neural mechanisms underlying the behavioral transition from pup-directed aggression to paternal care. Functional evidence suggested that this shift may be mediated by dynamic changes within Esr1^MPOA^ neuronal populations. To explore this, we first performed *in vivo* calcium imaging to monitor Esr1^MPOA^ neuronal activity as hostile virgin males transitioned into sires. For this we injected AAV-DIO-GCaMP6m into the MPOA of adult Esr1-Cre virgin male mice to enable *in vivo* monitoring of Esr1^MPOA^ neuronal activity in behaving mice (Fig. 5a). Post hoc immunofluorescence analysis showed that over 90% of GCaMP6m-labeled neurons co-expressed Esr1, confirming that the recorded calcium signals predominantly from Esr1^MPOA^ neurons (Fig. 5a-5c). We found that 4 out of 8 virgin male mice exhibited pup-directed aggression. Specifically, upon first exposure to pups, these hostile virgin males showed a transient and modest increase in Esr1^MPOA^ calcium activity, primarily during pup approach and investigation, with little to no response observed during attacks (Fig. 5d-5e). The pup was promptly removed from the cage after recording to prevent further physical attack toward the pups. Subsequently, these virgin males were paired with WT females and allowed to become sires. Remarkably, these sires exhibited a robust increase in Esr1^MPOA^ calcium activity beginning at pup approach, rising through the investigation, and peaking during retrieval, with elevated activity sustained throughout the retrieval phase (Fig. 5d-5g). These findings indicate that Esr1^MPOA^ neurons display significantly stronger and more sustained responses to pup stimuli in sires compared to hostile virgin males. In addition, Esr1^MPOA^ neurons were also strongly activated during the initial encounters with a female intruder or male intruder; however, these neurons showed no significant increase in activity when exposed to basic objects, a fake pup, or water spray (Extended Data Fig. 10a-10c). Importantly, no fluorescence changes were detected in the control 405-nm isosbestic channel, confirming that the observed signals reflected calcium-dependent activity (Extended Data Fig. 11a-11c; Extended Data Fig. 12a-12c).

**Figure 5.**
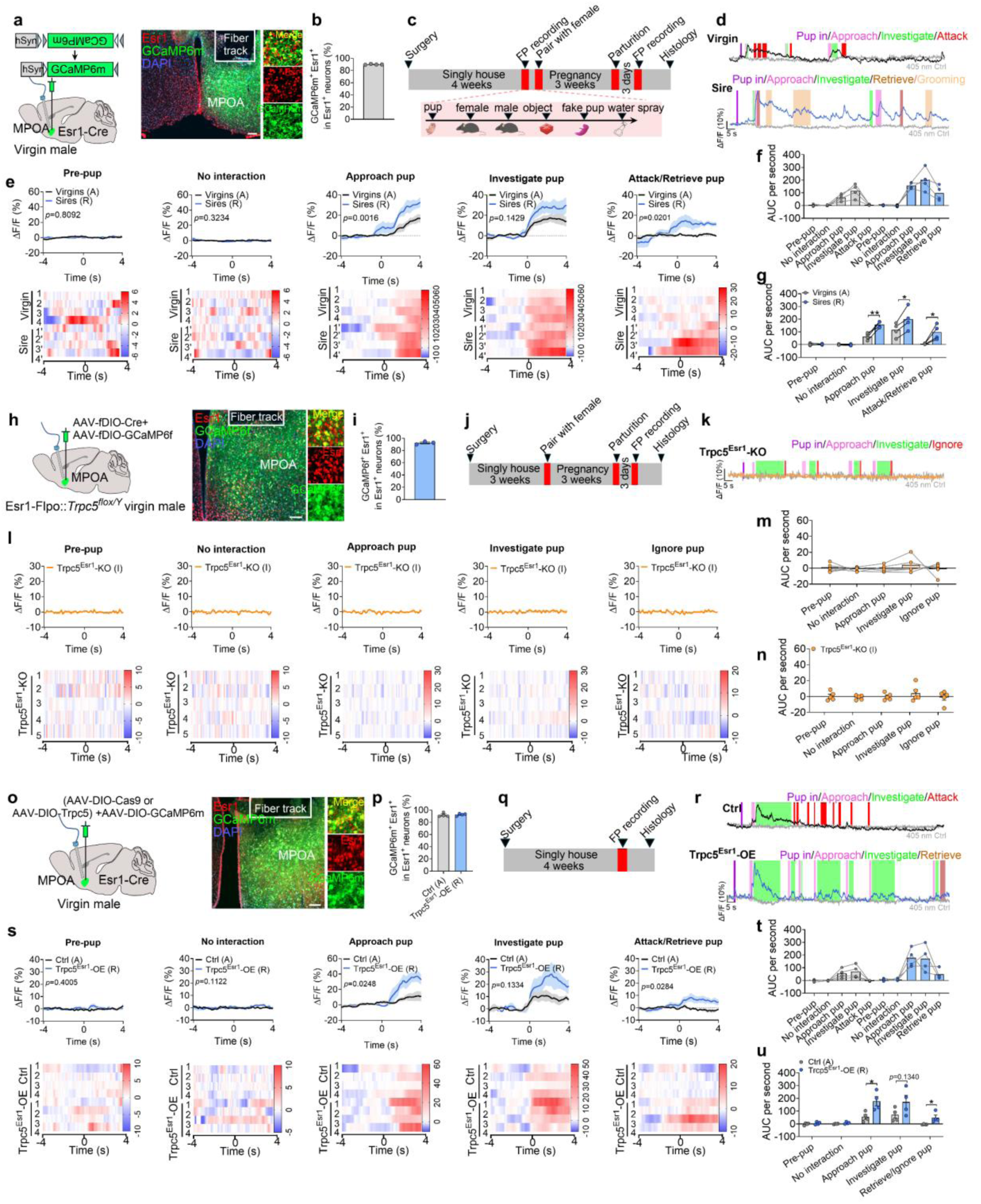
Trpc5 regulates Esr1^MPOA^ neuronal responses during the transition to fatherhood. (a) Left: schematic of the fiber photometry recording setup targeting Esr1^MPOA^ neurons. Right: representative image from four mice showing fiber track placement in the MPOA (white line) and the overlap between Esr1 immunostaining (red) and GCaMP6m expression (green). (b) Quantification of the percentage of GCaMP6m-expressing cells that co-express Esr1 in the MPOA (n = 4 mice, 16 weeks of age). Scale bars, 100 μm. (c) Experimental timeline and schematic overview of behavioral testing procedures. (d) Representative ΔF/F calcium traces of Esr1^MPOA^ neurons during pup interaction in a hostile virgin male (top) and a sire (bottom). (e) The ΔF/F calcium signal (top) and heatmap (bottom) of Esr1^MPOA^ neurons aligned to the onset of the following behaviors: pre-pup, no interaction, approach pup, investigate pup and attack/retrieve pup (from left to right). (f-g) The mean AUC of the ΔF/F during various pup-directed behaviors to compare responses across behaviors in hostile virgin males and sires (f) and responses of the same behavior between hostile virgin males and sires (g) (n = 4 mice, 12 weeks of age). (h) Left: schematic of the fiber photometry recording setup targeting and recording Esr1^MPOA^ neurons in Trpc5^Esr1^-KO sires. Right: a representative image showing fiber track placement in the MPOA (white line) and the overlap between Esr1 immunostaining (red) and GCaMP6m expression (green). (i) Quantification of the percentage of GCaMP6f-expressing cells that co-express Esr1 in the MPOA in Trpc5^Esr1^-KO sires (n = 3 mice, 18 weeks of age). Scale bars, 100 μm. (j) Experimental timeline and schematic overview of behavioral testing procedures. (k) Representative ΔF/F calcium traces of Esr1^MPOA^ neurons during pup interaction in a Trpc5^Esr1^-KO sire (bottom). (l) The ΔF/F calcium signal (top) and heatmap (bottom) of Esr1^MPOA^ neurons aligned to the onset of the following behaviors: pre-pup, no interaction, approach pup, investigate pup and attack/retrieve pup (from left to right). (m-n) The mean AUC of the ΔF/F during various pup-directed behaviors to compare responses across behaviors in Trpc5^Esr1^-KO sires (m) and responses of the same behavior in Trpc5^Esr1^-KO sires (n) (n = 5 mice, 18 weeks of age). (o) Left: Schematic of the fiber photometry recording setup targeting and recording Esr1^MPOA^ neurons in hostile virgin males and Trpc5^Esr1^-OE virgin males. Right: a representative image showing fiber track placement in the MPOA (white line) and the overlap between Esr1 immunostaining (red) and GCaMP6m expression (green). (p) Quantification of the percentage of GCaMP6f-expressing cells that co-express Esr1 in the MPOA between in hostile virgin males and Trpc5^Esr1^-OE virgin males (n = 4 mice per group, 14 weeks of age). Scale bars, 100 μm. (q) Experimental timeline and schematic overview of behavioral testing procedures. (r) Representative ΔF/F calcium traces of Esr1^MPOA^ neurons during pup interaction in a in hostile virgin males (top) and Trpc5^Esr1^-OE virgin males (bottom). (s) The ΔF/F calcium signal (top) and heatmap (bottom) of Esr1^MPOA^ neurons aligned to the onset of the following behaviors: pre-pup, no interaction, approach pup, investigate pup and attack/retrieve pup (from left to right). (t-u) The mean AUC of the ΔF/F during various pup-directed behaviors to compare responses across behaviors in hostile virgin males and Trpc5^Esr1^-OE virgin males (t) and responses of the same behavior in hostile virgin males and Trpc5^Esr1^-OE virgin males (u) (n = 4 mice, 12 weeks of age). Data presented as mean ± SEM, p value determined using two-way ANOVA (f,m,r), paired t-test (g), unpaired t-test (n,u). *p < 0.05, ***p < 0.001 and ****p < 0.0001.

To determine whether Trpc5 is required for pup-induced activation of Esr1^MPOA^ neurons during fatherhood, we injected a mixture of AAV-fDIO-GCaMP6f and AAV-fDIO-Cre into the MPOA of Esr1-Flpo::Trpc5^flox/Y^ mice (Fig. 5h-5j), and recorded calcium activity in the sires. Notably, deletion of Trpc5 from Esr1^MPOA^ neurons attenuated neuronal activation during pup-directed behaviors, including approach, investigation, and retrieval (Fig. 5k-5n). These mice also showed minimal responses to both male and female intruders, objects, a fake pup, or water spray (Extended Data Fig. 10d-10f), and no fluorescence changes were detected in the control 405-nm isosbestic channel (Extended Data Fig. 11d-11f; Extended Data Fig. 12d-12f).

Conversely, overexpression of Trpc5 in Esr1^MPOA^ neurons was sufficient to trigger pup retrieval behavior, and Trpc5^Esr1^-OE virgin males displayed increased Esr1^MPOA^ calcium activity during pup approach, investigation, and retrieval, similar to natural sires (Fig. 5o-5s). Quantitative analyses confirmed elevated neural responses and paternal behavior sequences in Trpc5^Esr1^-OE virgin males, compared to hostile virgin males (Fig. 5t-5u). However, Trpc5^Esr1^-OE mice did not exhibit significant changes in Esr1^MPOA^ calcium activity in response to female intruders, male intruders, objects, a fake pup, or water spray (Extended Data Fig. 10g-10i), indicating that Trpc5 overexpression selectively enhances Esr1^MPOA^ in response to pup stimuli but not to other cues. Again, signals from the 405-nm isosbestic control channel were comparable between hostile virgin males and overexpression virgin males that showed retrieval (Extended Data Fig. 11g-11i; Extended Data Fig. 12g-12i). Together, these results demonstrate that Trpc5 is necessary and sufficient to modulate Esr1^MPOA^ neuronal responses to pup stimuli, and is a critical molecular regulator of male paternal behavior.

### Trpc5 modulates the intrinsic excitability of Esr1^MPOA^ neurons

We next investigated the physiological mechanism underlying the enhanced *in vivo* responses observed in Esr1^MPOA^ neurons earlier. Towards this, we performed *in vitro* whole-cell current-clamp recordings from Esr1^MPOA^ neurons in hostile virgin males and sires (Fig. 6a). We observed clear state-dependent differences in neuronal excitability. While neurons from both groups increased their spiking activity in response to current injections, Esr1^MPOA^ neurons in sires exhibited significantly greater excitability, as evidenced by spike frequency-current (F-I) curves (Fig. 6b-6c), and a higher maximal number of action potentials (Fig. 6d) compared to hostile virgin males with increasing current. In addition, Esr1^MPOA^ neurons from sires displayed elevated baseline firing frequencies and more depolarized resting membrane potentials compared to those from hostile virgin males (Fig. 6e-6f). Together, these results support that fatherhood enhances the intrinsic excitability of Esr1^MPOA^ neurons, potentially facilitating their heightened responsiveness to pup-related stimuli *in vivo*.

**Figure 6.**
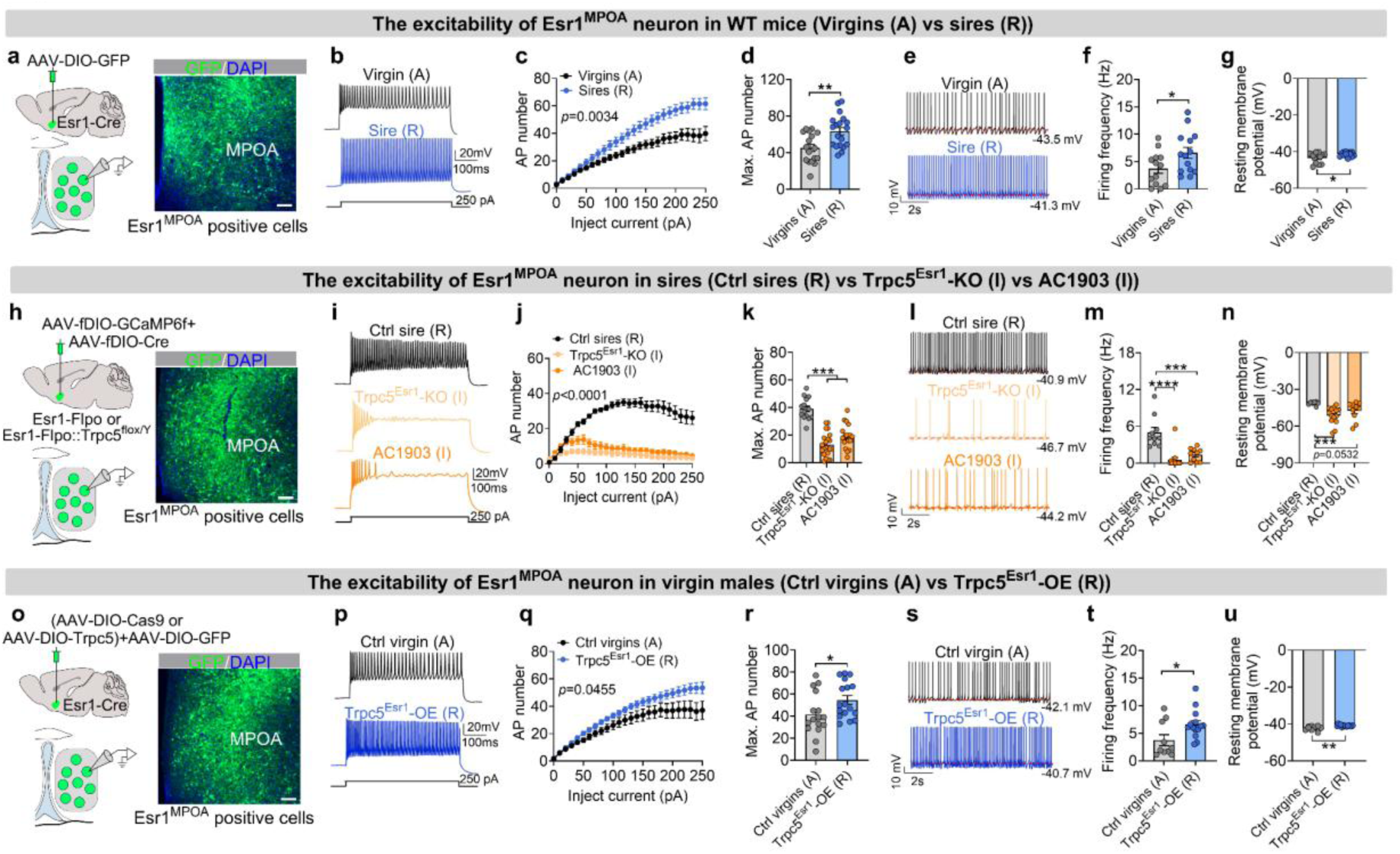
Trpc5 regulates excitability of Esr1^MPOA^ neurons during the transition to fatherhood. (a) Left: schematic of Cre-dependent AAV vectors used to express GFP in Esr1^MPOA^ neurons; Right: a representative image showing GFP-labeled Esr1^MPOA^ neurons in hostile virgin males and pup-retrieving sires, respectively. Scale bars, 100 μm. (b) Representative recording traces firing rates in response to depolarizing current injections of Esr1^MPOA^ neurons from hostile virgin males and pup-retrieving sire. (c) Frequency-current (F-I) curves of Esr1^MPOA^ neurons from hostile virgin male (17 neurons from 3 mice, 14 weeks of age) and pup-retrieving sires (19 neurons from 3 mice, 14 weeks of age). (d) The maximum action potential number of Esr1^MPOA^ neurons from hostile virgin males (17 neurons from 3 mice, 14 weeks of age) and pup-retrieving sires (19 neurons from 3 mice, 14 weeks of age) with maximally 250 pA injected current. (e) Representative recording traces of spontaneous firing at basic state from hostile virgin males and pup-retrieving sire. (f-g) Quantification of firing frequency (f) and resting membrane potential (g) in Esr1^MPOA^ neurons from hostile virgin males (13 neurons from 3 mice, 15 weeks of age) and pup-retrieving sires (14 neurons from 3 mice, 15 weeks of age). (h) Left: schematic of Flpo-dependent AAV vectors used to express Cre recombinase and GFP in Esr1^MPOA^ neurons from Esr1-Flpo and Esr1-Flpo::Trpc5^flox/Y^ male mice; Right: a representative image showing GFP-labeled Esr1^MPOA^ neurons, enabling targeted recordings of Esr1^MPOA^ neurons with or without Trpc5 deletion in sires. (i) Representative recording traces firing rates in response to depolarizing current injections of Esr1^MPOA^ neurons from control sire, Trpc5^Esr1^-KO sire, and control sire with AC1903 infusion in response to depolarizing current injections. (j) Frequency-current (F-I) curves of Esr1^MPOA^ neurons from control sires (15 neurons from 3 mice, 19 weeks of age), Trpc5^Esr1^-KO sires (22 neurons from 3 mice, 19 weeks of age), and control sires with AC1903 infusion (18 neurons from 3 mice, 19 weeks of age). (k) The maximum action potential number of Esr1^MPOA^ neurons from control sires (15 neurons from 3 mice, 19 weeks of age), Trpc5^Esr1^-KO sires (22 neurons from 3 mice, 19 weeks of age), and control sires with AC1903 infusion (18 neurons from 3 mice, 19 weeks of age) with maximally 250 pA injected current. (l) Representative recording traces of spontaneous firing at basic state from control sire, Trpc5^Esr1^-KO sire, and control sire with AC1903 infusion. (m-n) Quantification of firing frequency (m) and resting membrane potential (n) in Esr1^MPOA^ neurons from control sires (10 neurons from 3 mice, 19 weeks of age), Trpc5^Esr1^-KO sires (14 neurons from 3 mice, 19 weeks of age), and control sires with AC1903 infusion (10 neurons from 3 mice, 19 weeks of age). (o) Left: schematic of Cre-dependent AAV vectors used to overexpress Trpc5 and GFP in Esr1^MPOA^ neurons from Esr1-Cre male mice; Right: representative image showing GFP-labeled Esr1^MPOA^ neurons, enabling targeted recordings of Esr1^MPOA^ neurons with or without Trpc5 overexpression in virgin males. Scale bars, 100 μm. (p) Representative traces of action potential firing in Esr1^MPOA^ neurons from virgin males with or without Trpc5 overexpression in response to depolarizing current injections. (q) Frequency-current (F-I) curves of Esr1^MPOA^ neurons from virgin males with (17 neurons from 3 mice, 14 weeks of age) or without (17 neurons from 3 mice, 14 weeks of age) Trpc5 overexpression. (r) The maximum action potential number of Esr1^MPOA^ neurons from virgin males with (17 neurons from 3 mice, 14 weeks of age) or without (17 neurons from 3 mice, 14 weeks of age) Trpc5 overexpression with maximally 250 pA injected current. (s) Representative recording traces of spontaneous firing at basic state from virgin males with or without Trpc5 overexpression. (t-u) Quantification of firing frequency (t) and resting membrane potential (u) in Esr1^MPOA^ neurons from hostile virgin males (10 neurons from 3 mice, 14 weeks of age) and sires (14 neurons from 3 mice, 14 weeks of age). Data presented as mean ± SEM, p value determined using two-way ANOVA (c,j,q), unpaired t-test (d,f,g,r,t,u) and one-way ANOVA (k,m,n). *p < 0.05, ***p < 0.001 and ***p < 0.001.

To determine whether the increased excitability of Esr1^MPOA^ neurons in sires requires Trpc5 expression, we performed whole-cell recordings from Esr1^MPOA^ neurons in WT sires, Trpc5^Esr1^-KO sires, and WT sires treated with AC1903, the selective Trpc5 inhibitor (Fig. 6h). Both genetic deletion of Trpc5 and pharmacological inhibition with AC1903 significantly reduced neuronal excitability, as evidenced by fewer action potentials and decreased maximal spike counts in response to current injection (Fig. 6i-6k). Additionally, neurons from Trpc5^Esr1^-KO and AC1903-treated sires showed reduced firing frequency and exhibited slightly but significantly more hyperpolarized resting membrane potentials compared to WT sires (Fig. 6l-6n). Finally, we tested whether Trpc5 overexpression was sufficient to enhance the excitability of Esr1^MPOA^ neurons (Fig. 6o). In Trpc5^Esr1^-OE mice, Esr1^MPOA^ neurons exhibited significantly more action potentials in response to current injections (Fig. 6p-6q), along with increased maximal spike counts (Fig. 6r), firing frequencies, and depolarized resting membrane potentials (Fig. 6s-6u). Together, these findings demonstrate that the heightened excitability of Esr1^MPOA^ neurons during fatherhood is critically dependent on Trpc5 expression.

### Activation of Esr1^MPOA^ neurons reduces pup-directed aggression and increases paternal behavior

Our findings above led us to next hypothesize that activation of Esr1^MPOA^ neurons would (1) suppress inter-male aggression, and (2) trigger pup-retrieval behavior. To test this, we first used a chemogenetic approach to directly activate Esr1^MPOA^ neurons. To this end, we bilaterally injected a Cre-dependent excitatory DREADD (hM3Dq-mCherry) into the MPOA of Esr1-Cre virgin male mice (Fig. 7a). Electrophysiological recordings confirmed that hM3Dq-expressing neurons became depolarized and exhibited increased firing rates in response to clozapine-N-oxide (CNO) (Fig. 7b-7e). Consistently, *in vivo* CNO administration induced a greater than five-fold increase in c-Fos expression within Esr1^MPOA^ neurons (Extended Data Fig. 13a), indicating robust neuronal activation. Although activation of Esr1^MPOA^ neurons led to a progressive decrease in body temperature over time (Extended Data Fig. 13b), the locomotor activity was unaffected by CNO injection after 30 minutes (Extended Data Fig. 13c-13d). Based on this observation, we selected the 30-minute time point as the paternal behavioral testing window during retrieval assay. During this time period, CNO-mediated activation of Esr1^MPOA^ neurons significantly reduced the proportion of virgin males that displayed pup-directed aggression compared to saline-injected controls (Fig. 7f-7g). Moreover, both the latency to attack and the cumulative probability of attack were markedly reduced following neuronal activation (Fig. 7h-7i). Moreover, activation of Esr1^MPOA^ neurons did not induce pup retrieval in virgin males but showed a significant increase in pup investigation relative to saline injection (Fig. 7j-7k). These behaviors were not observed in control mice, regardless of CNO or saline treatment.

**Figure 7.**
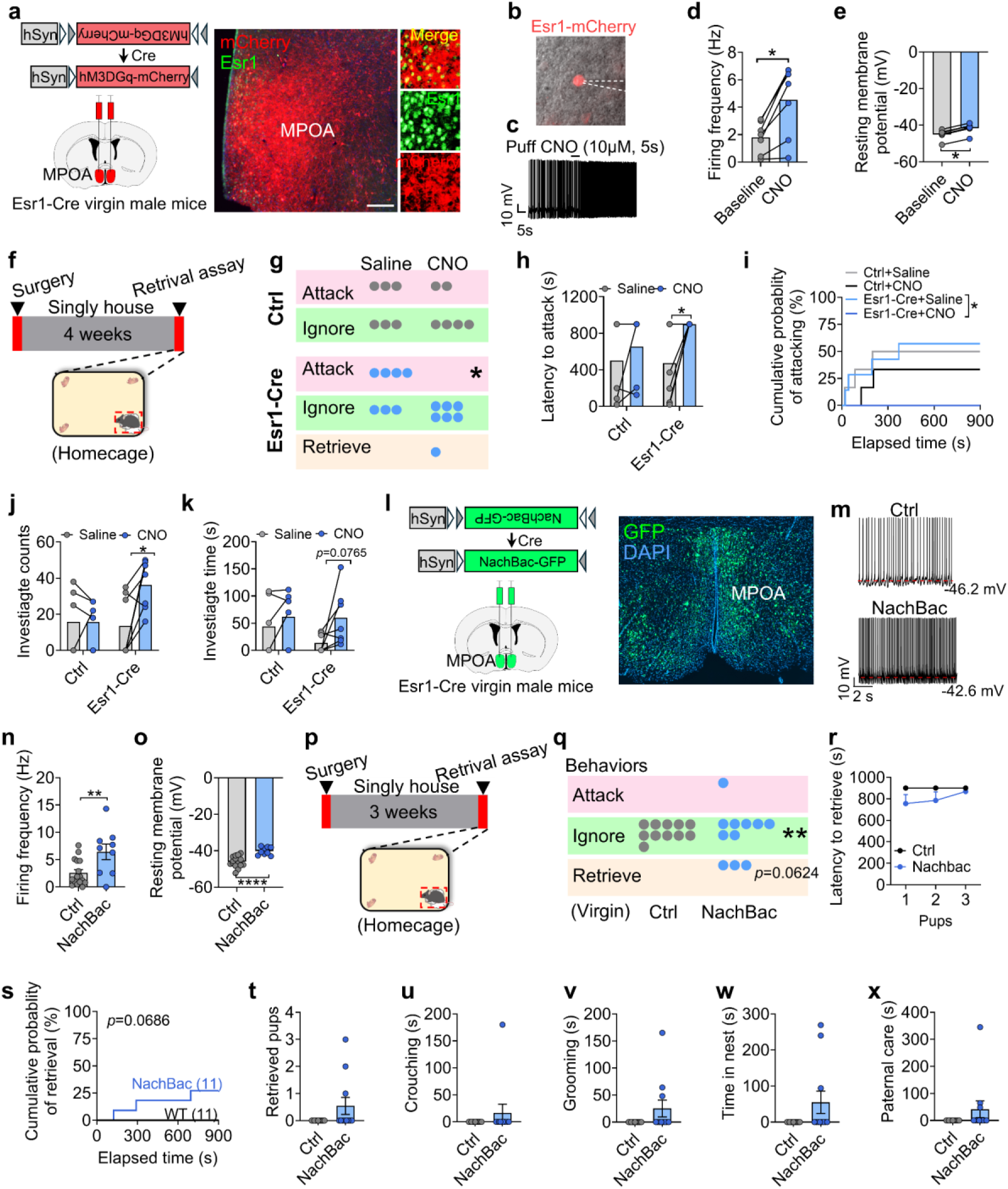
Acute activation of Esr1^MPOA^ neurons reduces pup-directed aggression, while chronic activation enhances paternal behavior. (a) Left: schematic of injection of Cre-dependent excitatory DREADD (hM3Dq-mCherry) into the MPOA of Esr1-Cre virgin male mice; Right: a representative image showing the mCherry (red) expression in Esr1^MPOA^ positive neurons (green). (b) A hM3Dq-mCherry expressing neuron in MPOA targeted for patch-clamp recording. (c) Example of a patched hM3Dq-mCherry labeled neuron subjected to a CNO puff (10 μM, 5 s). (d-e) Electrophysiological recordings revealed that hM3Dq-expressing neurons exhibited increased firing frequency (d) and membrane depolarization (e) in response to CNO. (f) Experimental timeline. (g) Dots indicate the number of control and Esr1-Cre virgin males treated with saline or CNO that exhibited attack, ignore, or retrieve. (h) Latency to attack the pups (n= 6-7 mice per group, 12 weeks of age). (i) Cumulative probability of pup attacking (n= 6-7 mice per group, 12 weeks of age). (j-k) The quantification of pup investigation counts (j) and duration (k) in control and Esr1-Cre virgin males treated with saline or CNO (n = 6-7 mice per group, 12 weeks of age). (l) Left: schematic of injection of Cre-dependent NachBac-GFP into the MPOA of control and Esr1-Cre virgin male mice; Right: a representative image showing the GFP expression in MPOA. (m) Representative traces showing typical AP firing in neurons recorded from the control (top) and NachBac-mediated AP in neurons (bottom). (n-o) The firing frequency (n) and resting membrane potential (o) from control and NachBac group. (p) Experimental timeline. (q) Dots indicate the number of control and Esr1-Cre virgin males expressing NachBac that exhibited attacking, ignoring, or retrieving (n= 11 mice per group, 11 weeks of age). (r) The latency time to retrieve pups (n= 11 mice per group, 11 weeks of age). (s) Cumulative probability of pup retrieval (n= 11 mice per group, 11 weeks of age). (t) The retrieved pups’ numbers (n= 11 mice per group, 11 weeks of age). (u-x) The duration of crouching pups (u), grooming pups (v), time spent in the nest (w) and paternal care (x) (n= 11 mice per group, 11 weeks of age). Data presented as mean ± SEM. p value determined using two-way ANOVA (h,j,k,r), Mann-Whitney test (t,u,v,w,x), chi-squared test (g,q) and paired t test (d,e). *p < 0.05, **p < 0.01 and ****p < 0.0001.

Next, we injected a Cre-dependent AAV vector expressing a bacterial sodium channel (AAV-EF1a-Flex-EGFP-P2A-mNachBac-GFP, referred to as AAV-DIO-NachBac-GFP) into the MPOA of Esr1-Cre virgin male mice to drive chronic activation of Esr1^MPOA^ neurons (Fig. 7l). WT virgin male mice received the same virus injections and served as controls. To first validate functional modification of these neurons, we performed electrophysiological recordings, which revealed an increased spontaneous firing and a decreased resting membrane potential (Fig. 7m-7o). Also, we unilaterally injected AAV-DIO-NachBac-GFP into the right side of MPOA and observed robust c-Fos expression in NachBac-expressing Esr1^MPOA^ neurons compared to the un-injected side, indicating sustained neuronal activation using this viral tool (Extended Data Fig. 14a-14b). Behaviorally, chronic activation of Esr1^MPOA^ neurons significantly reduced pup-ignoring behavior and modestly promoted pup retrieval in virgin male mice (Fig. 7p-7r), as evidenced by an increased cumulative retrieval probability (Fig. 7s). However, it did not affect the number of pups retrieved, or other related parameters (Fig. 7t-7x). Also, activation of these neurons did not affect general locomotor activity, anxiety-like behaviors in open field tests (Extended Data Fig. 14c-14g), or depression-related behaviors in forced swim (Extended Data Fig. 14h) and tail suspension tests (Extended Data Fig. 14i). Social interaction ability also remained unchanged after chronic activation of Esr1^MPOA^ neurons (Extended Data Fig. 14j-14o). However, activation of Esr1^MPOA^ neurons significantly reduced inter-male aggression toward both adult (Extended Data Fig. 14p-14r) and juvenile intruders (Extended Data Fig. 14s-14u), consistent with the suppression of pup-directed aggression.

### Trpc5 overexpression enhances escape behavior and elicits exploratory diving behavior **in** virgin male mice

Interestingly, exposure to predator odor (TMT) significantly suppressed locomotor activity and increased freezing behavior in control but not Trpc5^Esr1^-OE mice (Fig. 8a-8e). Further, we employed a multi-phase escape task with progressively increasing difficulty to assess the escaping capability^26^. Trpc5^Esr1^-OE mice demonstrated significantly reduced latency to escape and higher escape success rates across phases without any prior training or habituation, reflecting enhanced escaping capability (Fig. 8f-8i). Strikingly, we found that Trpc5^Esr1^-OE mice showed a marked increase in voluntary diving counts and duration in a 12 cm-deep water bath at either 25°C or 30°C temperature (Fig. 8j-8l; Supplementary Movie 1). To further examine this behavior, we tested them in larger tanks with water depths of 18 cm or 28 cm using 25°C water temperature (Fig. 8m). Remarkably, Trpc5^Esr1^-OE mice continued to dive voluntarily, often reaching the bottom multiple times (Fig. 8n-8o), potentially reflecting an enhanced exploratory response. Together, these results demonstrate that Trpc5 overexpression in Esr1^MPOA^ neurons amplifies a broad spectrum of defensive behaviors, with particularly robust effects on exploratory behavior/diving and escaping capabilities, highlighting a potential role of Trpc5 in modulating behavioral reactivity to environmental threats. Interestingly, NachBac-induced activation of Esr1^MPOA^ did not elicit diving behavior, although the escaping capability was enhanced in these mice (Extended Data Fig. 15a-15k).

**Figure 8.**
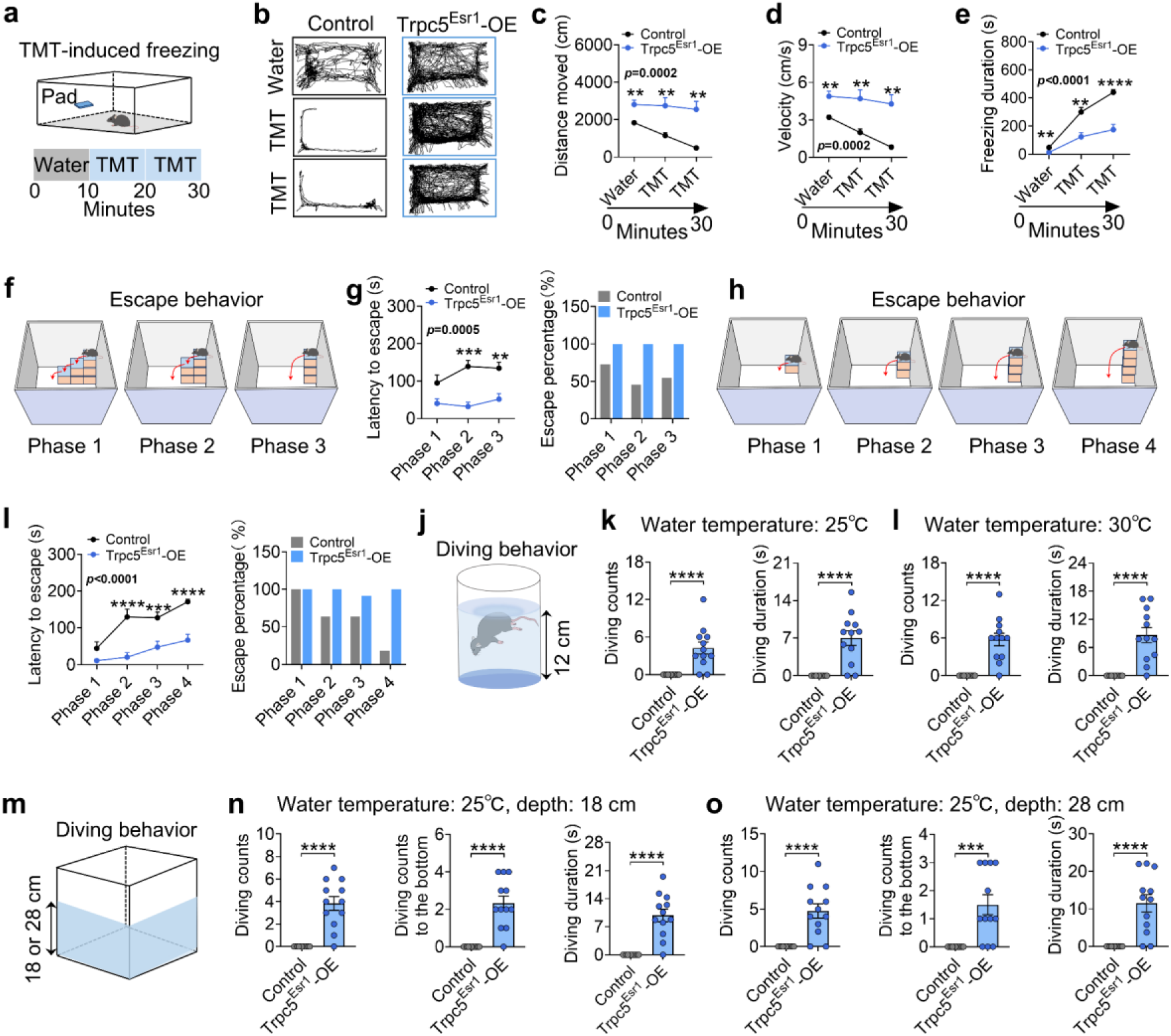
Trpc5 overexpression in Esr1^MPOA^ neurons enhances escape behavior and elicits exploratory diving behavior. (a) Illustration showing TMT-induced behavior in the home cage. Water was first added to the pad for 10 minutes, followed by replacement with a new pad containing TMT for an additional 20 minutes. (b) The track of movement. (c-d) The distance moved (c) and velocity (d) duration within 10 minutes across the water, TMT, and TMT exposure (n = 11-12 mice per group, 18 weeks of age). (e) Freezing duration within 10 minutes across the water, TMT and TMT exposure. (n = 11-12 mice per group; 17 weeks of age). (f) Schematic illustrating a multi-phase escape task with progressively increasing difficulty from phase 1 to 3. (g) Quantification of escape latency (left) and percentage of mice that successfully escaped (right) in control and Trpc5^Esr1^-OE virgin male mice (n = 11-12 mice per group; 16 weeks of age). (h) Schematic illustrating a multi-phase escape task with progressively increasing difficulty from phase 1 to 4. (i) Quantification of escape latency (left) and percentage of mice that successfully escaped (right) in control and Trpc5^Esr1^-OE virgin male mice (n = 11-12 mice per group; 16 weeks of age). (j) Illustration showing diving behavior in a glass beaker (beaker height: 18 cm; water depth: 12 cm). (k-l) Diving counts and durations in control and Trpc5^Esr1^-OE virgin male mice tested in 25 °C (k) or 30 °C (l) water (n = 11-12 mice per group, 15 weeks of age). (m) Illustration showing diving behavior in an open field arena (arena length x width x depth: 42 x 42 x 30 cm; water depth: 18 or 28 cm). (n) Diving counts (left), diving counts to the bottom (middle) and diving durations (right) in control and Trpc5^Esr1^-OE virgin male mice tested at 18 cm water depth (n = 11-12 mice per group, 15 weeks of age). (o) Diving counts (left), diving counts to the bottom (middle) and diving duration (right) in control and Trpc5^Esr1^-OE virgin male mice tested at 28 cm water depth (n = 11-12 mice per group, 15 weeks of age). Data presented as mean ± SEM, p value determined using two-way ANOVA (c,d,e,j,i), and Mann-Whitney test (k,l,n,o). *p < 0.05, **p < 0.01, ***p < 0.001 and ****p < 0.0001.

## Discussion

Here we demonstrate that male mice carrying a human TRPC5 loss-of-function mutation exhibit impaired paternal behaviors. Interestingly, the behavioral shift from infanticide in virgin males to caregiving in wild-type sires is associated with an upregulation of Trpc5 expression in Esr1^MPOA^ neurons. While selective deletion of Trpc5 from Esr1^MPOA^ neurons diminishes paternal behaviors in sires, remarkably, selective overexpression of Trpc5 in these neurons converts otherwise infanticidal virgin males to exhibit paternal care just like sires. Further, we show that excitability of Esr1^MPOA^ neurons undergoes experience-dependent changes during this transition, critically relying on Trpc5 expression. Notably, Trpc5 overexpression also triggers robust escape and exploratory behaviors linked to survival-seeking responses. Collectively, these findings highlight a pivotal role for Trpc5 in Esr1^MPOA^ neurons in parental behavior.

Parental care is essential for the survival of vulnerable offspring and the continuation of the species. However, parental behavior is highly sex-dependent, with males and females showing distinct infant-directed behaviors. Virgin females typically avoid pups or show weak maternal responses, which increase markedly during pregnancy and lactation. In contrast, sexually naïve males often attack offspring, whereas experienced sires retrieve pups. These observations indicate that both sex and prior parenting experience shape sensitivity to offspring cues and caregiving efficiency in mice^27^. Decades of research have established the critical roles of maternal behaviors, such as retrieving, grooming, and nursing, in promoting reproductive success and fostering prosocial development^28–31^. Over the past several decades, the biological mechanisms underlying direct paternal care in male animals have received increasing attention^32–34^. Hormonal regulation plays a central role in maternal behavioral transitions. Lactogenic hormones (prolactin, estradiol, and progesterone) act through receptors in key brain regions to facilitate maternal responses in dams^25,30^. By contrast, males lack such endocrine cycles, raising the question of how paternal behaviors arise without classic hormonal cues^17^. Previously, we found that Trpc5 expression in PVH neurons is essential for dams to express maternal behavior^23^. However, PVH Trpc5 expression levels are comparable between hostile virgin males and sires, suggesting a different mechanism. Here we show that an upregulation of Trpc5 expression in Esr1^MPOA^ neurons is associated with the transition from infanticidal virgin males to caregiving sires, and that deletion of Trpc5 from these neurons diminishes paternal behaviors in sires. Thus, we suggest that Trpc5 signaling in the brain regulates both maternal and paternal behaviors through distinct neural substrates.

In Trpc5-overexpressing mice, behaviors such as enhanced escaping and exploratory diving suggest that survival-oriented strategies are integral to paternal adaptations^26,35–37^. These behaviors promote rapid environmental assessment, resource acquisition, and predator avoidance, thereby protecting offspring through enhanced sire survival. From an evolutionary perspective, such traits may have been co-opted to support paternal care. Notably, we report for the first time a voluntary diving phenotype in water-based tests with no prior training needed, potentially reflecting a conserved exploratory strategy in hostile environments. Its emergence with Trpc5 overexpression suggests a broader role for this channel in orchestrating context-specific adaptive responses. Future work should elucidate the physiological mechanisms underlying these behaviors and their contributions to survival.

Trpc5 is a calcium-permeable cation channel highly expressed in the central nervous system ^22,38^. Mechanistically, Trpc5 plays dual roles in paternal behavior: it elevates the baseline activity of Esr1^MPOA^ neurons, and enhances their responsiveness to pup-related stimuli. Even in virgin males, Esr1^MPOA^ neurons exhibit some activation, but fatherhood induces TRPC5-dependent remodeling that markedly amplifies neuronal excitability and drives the behavioral shift from infanticide to caregiving. Through transducing sensory and internal cues into electrical signals, Trpc5 enables Esr1^MPOA^ neurons to integrate environmental information with internal states, providing a mechanistic basis for both Step I (suppression of infanticide) and Step II (initiation of caregiving) in the two-step model of paternal behavior^11^. Importantly, DREADD or NachBac-mediated activations increase Esr1^MPOA^ firing but fail to fully recapitulate the behavioral outcomes of Trpc5 overexpression, underscoring that Trpc5 is not a mere excitability enhancer. Beyond regulating neuronal firing as a receptor-activated calcium-permeable channel, Trpc5 functions as a versatile molecular hub, coordinating protein-protein interactions, scaffolding, and intracellular signaling. For example, Trpc5 forms homo- and heteromeric assemblies with Trpc1, Trpc3, and Trpc4, regulating dendritic spine morphology, synaptic plasticity, and receptor clustering. In hippocampal neurons, Trpc5-Trpc1 heteromers modulate spine structure and plasticity, illustrating how heteromerization broadens gating and receptor coupling^39^. Further, Trpc5 can interact with calmodulin and calcium binding protein 1 to convert transient calcium influx into sustained intracellular signals^40,41^. Thus, multiple molecular/cellular mechanisms, including channel activity, protein interactions, and fine-tune neuronal excitability and circuit integration, can contribute to the complex behaviors mediated through Trpc5, including the complete behavioral spectrum of paternal care.

The developmental transition to motherhood requires gene expression changes that remodel the brain, enabling females to perform maternal behaviors. In postpartum female mice, gene expression undergoes significant alterations, affecting neural activity, synaptic plasticity, and hormone signaling pathways^42^. Motherhood is associated with a pronounced increase in calcitonin receptor expression, which likely contributes to enhanced synaptic plasticity in the MPOA, particularly during the early postpartum period^15^. In contrast, only a few studies, mostly in fish, have documented dramatic changes in gene expression as males become fathers^43^. Fatherhood is also associated with a relative decrease in overall MPOA gene expression related to synaptic transmission and plasticity, reflecting sex-specific and experience-dependent adaptations^44^. A key mechanism underlying brain plasticity involves modifications in the strength of synaptic connections, such as those observed in PVH oxytocin neurons and the lateral hypothalamic area^11^. Interestingly, Trpc5 is upregulated in sires, suggesting that while global synaptic plasticity may decline, specific molecular pathways are selectively enhanced to support paternal behavior, increase neuronal excitability, and heighten responsiveness to offspring-related cues. The mechanisms underlying the increased Trpc5 expression in sires remain unclear. The increases in both mRNA and protein levels suggest a mechanism at least partially involving enhanced Trpc5 gene transcription, although the detailed transcriptional control, especially during the development of paternal behavior, warrant future investigations.

Together, our study identifies Trpc5 as a critical molecular mediator of paternal behavioral plasticity, enabling the transition from infanticidal to caregiving responses in males. These findings broaden the definition of fatherhood to include both direct caregiving and adaptive survival behaviors that indirectly enhance offspring protection. Ultimately, this work provides a framework for exploring how gene-specific plasticity supports the emergence of paternal care and its evolutionary significance.

## Methods and Materials

### Mice

All mouse lines were maintained on a C57BL/6J background. Adult male mice (8-20 weeks old) were used in all experiments. For the Trpc5^K34del^ mutant line, male Trpc5^K34del/Y^ mice and their wild-type (WT) littermates were generated and used following the same protocol^23^. Esr1-Cre (#017911) and Esr1-Flpo (#037009) mice were obtained from the Jackson Laboratory, and Trpc5^flox/flox^ mice were generously provided by Dr. Kevin Williams (UT Southwestern). Male Esr1-Flpo mice were crossed with female Trpc5^flox/flox^ mice to generate Esr1-Flpo::Trpc5^flox/Y^ male mice. WT C57BL/6J pups at postnatal days 1-3 (P1-P3) were used for pup retrieval behavioral experiments. Group-housed WT adult (2-4 months of age) and juvenile (21-28 days of age) were used as the intruders in resident-intruder assay. All animal procedures and care were approved by the Institutional Animal Care and Use Committee (IACUC) of Baylor College of Medicine. Mice were housed under controlled environmental conditions: temperature (22-24°C), humidity (60%), and a 12-hour light/dark cycle (lights off from 18:00 to 6:00). Food and water were provided ad libitum.

### Viruses

The following AAV vectors were obtained from Addgene: AAV9-hSyn-DIO-GCaMP6m.WPRE.SV40 (#100838; 2.7 × 10¹³ gc/ml), AAV1-Ef1a-fDIO-GCaMP6f (#128315; 1 × 10¹³ gc/ml), AAV9-hSyn-DIO-GFP (#100043; 4.3 × 10¹³ gc/ml), AAV9-hSyn-DIO-hM3D(Gq)-mCherry (#44361; 1 × 10¹³ gc/ml), and AAV9-EF1α-fDIO-Cre (#121675; 1 × 10¹³ gc/ml). AAV2-CMV7-DIO-saCas9 (#7122, 1 x 10^13^ gc/ml) was obtained from Vector Biolabs. AAV-DIO-NachBac-GFP and AAV-hSyn-DIO-Trpc5-Flag were generated from Baylor NeuroConnectivity Core. All viral preparations were aliquoted and stored at -80°C until use.

### Stereotaxic injection

Mice were anaesthetized with isoflurane (3-4%) and maintained under isoflurane (1-1.5%) throughout the surgery. The MPOA was targeted using the following coordinates relative to Bregma: anterior-posterior (AP) +0.02 mm, medial-lateral (ML) ±0.28 mm, and dorsal-ventral (DV) −5.20 mm.

To investigate the role of Trpc5 in Esr1^MPOA^ neurons, viral manipulations were performed in adult male mice. For deletion of Trpc5 in Esr1^MPOA^ neurons of sires, 200 nl of AAV9-EF1α-fDIO-Cre was bilaterally injected into the MPOA of adult Esr1-Flpo (control) or Esr1-Flpo::Trpc5^flox/Y^ mice. Three weeks post-surgery, males were paired with WT females and co-housed until three days after parturition. For Trpc5 overexpression in Esr1^MPOA^ neurons in virgin males, 400 nl of AAV-hSyn-DIO-Trpc5-Flag was bilaterally injected into the MPOA of adult WT and Esr1-Cre mice.

For in vitro recordings in virgins and sires, 200 nl of AAV9-hSyn-DIO-GFP was bilaterally injected into the MPOA of Esr1-Cre mice to label Esr1^MPOA^ neurons with green fluorescence; For in vivo recordings in virgin and sires, 200 nl of AAV9-hSyn-DIO-GCaMP6m was injected into the right MPOA of Esr1-Cre mice.

For the in vitro recording in the Trpc5 deletion study, both Esr1-Flpo and Esr1-Flpo::Trpc5^flox/Y^ mice bilaterally received a mixture of 300 nl AAV9-EF1α-fDIO-Cre and 200 nl AAV9-Ef1a-fDIO-GCaMP6f in the MPOA. For in vivo fiber photometry recording, Esr1-Flpo::Trpc5^flox/Y^ mice received 300 nl AAV9-EF1α-fDIO-Cre combined with 200 nl AAV1-Ef1a-fDIO-GCaMP6f in the right MPOA.

For the in vitro recording in Trpc5 overexpression study, Esr1-Cre mice bilaterally received a mixture of 300 nl AAV-DIO-Trpc5 and 200 nl AAV9-hSyn-DIO-GFP in the MPOA, and 300 nl AAV2-CMV7-DIO-saCas9 with 200 nl AAV9-hSyn-DIO-GFP in the MPOA as controls. For in vivo fiber photometry recording, Esr1-Cre mice received 300 nl AAV-DIO-Trpc5 in the left side of MPOA, and a mixture of 300 nl AAV-DIO-Trpc5 with 200 nl AAV9-hSyn-DIO-GCaMP6m in the right side of MPOA. As controls, Esr1-Cre mice received 300 nl AAV2-CMV7-DIO-saCas9 in the left side of MPOA, and a mixture of 300 nl AAV2-CMV7-DIO-saCas9 with 200 nl AAV9-hSyn-DIO-GCaMP6m in the right side of MPOA. For all fiber photometry recording studies, a Doric probe will be implanted 200 μm above the right side of MPOA, allowing in vivo recordings in freely behaving mice.

For chemogenetic activation of Esr1^MPOA^ neurons in virgin males, 200 nl of AAV9-hSyn-DIO-hM3D(Gq)-mCherry was bilaterally injected into the MPOA of adult WT and Esr1-Cre mice. For chronic activation of Esr1^MPOA^ neurons in virgin males, 200 nl of AAV-DIO-NachBac-GFP was bilaterally injected into the MPOA of adult WT and Esr1-Cre mice.

### Drug administration

CNO (#16882; Cayman) was first dissolved in DMSO and diluted in saline, then administered to mice via intraperitoneal (i.p.) injection at a dose of 0.5 mg kg^-1^. AC1903 (#1910; Tocris Bioscience) was first dissolved in DMSO, then diluted in a solution containing 5% Tween 80, 40% PEG 300, and 45% saline, and administered via i.p. injection at a final concentration of 50 mg kg^-1^.

### Mouse behavioral tests

#### Paternal behaviors assay for virgin males

For behavioral testing in virgin male mice, each test cage was supplied with shredded nestlet material, which the mouse typically used to construct a nest in one corner of the cage. The resident male’s cage and nest were left undisturbed for approximately one week before testing. During the assay, three WT C57BL/6J pups (postnatal day 1-3), which had not previously been exposed to any adult males, were placed in separate corners of the cage away from the resident male’s nest. The introduction of the pups marked the start of the behavioral test. Males were allowed to freely interact with the pups for 15 minutes. If any sign of pup-directed aggression was observed, the assay was terminated immediately, and the injured pup was euthanized. Each male was tested only once. Males were categorized into behavioral groups based on the following criteria: Retrieve: retrieved at least one pup to the nest; Attack: attacked at least one pup; Ignore: showed neither retrieval nor attack toward pups. Notably, animals that retrieved pups did not display aggression, and vice versa. Therefore, the sum of the proportions of these three behavioral categories (Attack, Ignore, Retrieve) always equaled 100%. In addition to these primary behavioral outcomes, the following behaviors were also scored: Latency to retrieve: time taken to pick up a pup and carry it to the nest; Grooming: licking of pups; Crouching: assuming a nursing-like posture over at least two pups; Nest building: collecting and organizing nesting material; Time spent in the nest. The total duration of grooming, crouching, and nest building was summed to define paternal care duration. If a male exhibited grooming while crouching, the time was counted toward both behaviors. For animals that either attacked or ignored pups during the 15-minute assay, the latency to retrieve was recorded as 15 minutes.

#### Paternal behaviors assay for sires

The behavioral assay for sires followed the same protocol used for virgin males, with specific modifications. Virgin males were individually housed and paired with a female. The following morning, the presence of a vaginal plug was checked; only males from pairs with a confirmed plug were included in subsequent experiments. Males remained cohabitated with the mated females until three days after parturition. The behavioral assay was conducted on postnatal day 3. Six hours prior to the test, the female and the relative litters were removed from the home cage, leaving only the sires. To assess paternal behavior, three unfamiliar pups (P1-P3), unrelated to the resident male and prepared as in the virgin male assay, were introduced into the cage. Behavioral scoring was performed as described for virgin males, except that the retrieval was defined when all three pups had been retrieved.

#### Forced swim test

The forced swim test (FST) was conducted as previously described^45^. Briefly, mice were placed into a glass beaker (height: 18 cm, diameter: 14 cm) filled with water at 23-25°C to a depth of 12 cm. Mice were allowed to swim for 6 minutes, and the immobility time during the last 4 minutes was quantified. At the end of the test, mice were gently removed from the beaker, blotted dry with a paper towel, and then returned to their home cage.

#### Tail suspension test

The tail suspension test (TST) was performed as previously described^45^. Each mouse’s tail was suspended 60 cm above the floor of the room, using tape placed less than 1 cm from the tip of the tail. Behavior was recorded for 6 minutes using a camera, and the last 4 minutes of the recording were analyzed to determine the total duration of immobility.

#### Open field test

The open field test (OFT) was performed in the open field arena (length × width × height: 42 × 42 × 30 cm). The lines divide the floor into sixteen evenly spaced squares (10.5 × 10.5 cm). The center consisted of four squares in the center of the device (21 × 21 cm). A mouse was first placed into the center of the area and allowed to explore for 5 minutes, and the paths of the animals were recorded by a video camera. Total distance travelled, velocity, distance travelled in the center, time spent in the center, and entries into the center were analyzed using the Noldus EthoVision XT (Noldus, Leesburg, VA, USA). In addition, the number of episodes of rearing and the total duration of rearing were recorded manually. The arena was cleaned with 75% alcohol solution between different mice.

#### Three-chamber social interaction test

The social interaction test (SIT) used a three-chambered box with openings between chambers for the mouse to pass through. The entire procedure consisted of a 5-minute habituation phase, followed by a 10-minute social interaction phase and a 10-minute social novelty phase. During the habituation, a subject mouse was placed in the behavior apparatus for 5 minutes with the chamber doors open. After the habituation session, the social preference test was performed. While the subject mouse was in the center chamber, a never-before-met intruder mouse A was placed in a pencil cup in the right chamber. Meanwhile, a pencil cup containing a novel object was placed in the left chamber. The subject mouse was allowed to move freely for 10 minutes, then the interaction time with each cup was manually recorded. Then, the novel object was replaced with another mouse B. The subject mouse was allowed to move freely for 10 minutes, then the interaction time with each cup was manually recorded again. The preference ratio was calculated by the following equations (OT: object time; M1T: mouse 1 time; M2T: mouse 2 time). Preference ratio in social interaction=(M1T−OT)/(M1T+OT)∗100%. Preference ratio in social novelty=(M2T−M1T)/(M2T+M1T)∗100%. Total distance travelled was analyzed using the Noldus EthoVision XT (Noldus, Leesburg, VA, USA).

#### Resident-intruder test

Mice were individually housed in their home cages for at least one weeks before behavioral testing. Cages were left unclean and unaltered to allow the accumulation of olfactory cues and to promote territorial behavior in the resident mouse. The assay began by introducing an unfamiliar adult male mouse (2-4 months of age) into the resident’s home cage. After 15 minutes of interaction, the intruder was removed. One week later, a second assay was conducted by introducing an unfamiliar juvenile male mouse (21-28 days of age) into the resident’s home cage for another 15-minute session. Throughout each 15-minute interaction, behavior was continuously recorded using a video camera. Videos were analyzed in a blind manner to assess the latency to first attack, total number of attacks, and cumulative time spent attacking by the resident mouse.

#### Diving behavior

To examine diving behavior in mice, we first placed them in a glass beaker (height: 18 cm, diameter: 14 cm) filled with water at 25 °C or 30 °C to a depth of 12 cm. One week later, the mice were exposed to a larger space-an open-field arena (42 × 42 × 30 cm)-filled with water at either 25 °C to a depth of 18 cm or 28 cm. The mice were gently placed into the water, and their diving behavior was recorded by a side-mounted camera. Diving counts and diving duration were subsequently analyzed.

#### TMT induced freezing behavior

In the new and clean home cage, a cotton pad was placed along the wall. Water (20 μl) was first added to the pad for 10 minutes, followed by replacement with a new pad containing TMT for an additional 20 minutes. Each mouse was individually introduced into the cage. The following parameters were analyzed: distance moved and total duration of freezing behavior.

#### Assessment of multi-phase escape behavior

In the empty box assay, mice were placed individually in a transparent open-field arena (42 × 42 × 30 cm) devoid of any objects or escape structures. The height of the walls (30 cm) was sufficient to prevent escape by jumping. In the downward escape assay, identical boxes (10 × 10 × 5 cm) were arranged in a staircase-like configuration. To successfully escape, animals were required to jump from the first to the final elevated box, which was positioned 5 cm above the floor, onto the ground. Each trial in the escapable assays lasted a maximum of 3 minutes (180 seconds). If a mouse failed to escape within this period, the trial was terminated, and escape latency was defined as 180 seconds. No habituation, familiarization, or prior training was provided before any of the behavioral assays.

#### Sexual behavior

Ten-week-old C57BL/6J virgin female mice, confirmed to be sexually receptive via vaginal smear, were placed into the cage of the resident male mice. Each interaction lasted 15 minutes, during which behavior was recorded on video and later analyzed for sniffing, latency to mount, mount counts, mount duration, intromission counts and duration.

#### Immunofluorescence

At the end of the study, mice were anesthetized with inhaled isoflurane and quickly perfused with saline, followed by 10% formalin. Brains were immediately collected and postfixed in ice-cold 5% formalin overnight, after which they were cryoprotected in 30% sucrose in PBS. Coronal brain sections (30 µm) were prepared and processed following a standard immunofluorescence protocol. Briefly, sections were washed three times in 0.1% PBST (PBS+0.1% Triton), with ten-minute intervals between washes. They were then blocked for 2 hours in 0.3% PBST (PBS+0.3% Triton) containing 5% goat normal serum.

To examine Trpc5 expression in Esr1^MPOA^ neurons, primary mouse anti-Trpc5 antibody (1:1000, 75-104, Antibodies Inc) was co-applied with either primary rabbit anti-Esr1 antibody (1:1000, 06-935, Millipore) to brain sections containing the MPOA from both male virgins and sires.

To examine Trpc5 expression in the PVH, primary mouse anti-Trpc5 antibody (1:1000, 75-104, Antibodies Inc) was applied to brain sections containing the PVH from both male virgins and sires.

To validate Trpc5 overexpression or deletion in Esr1^MPOA^ neurons, primary mouse anti-Trpc5 antibody (1:1000, 75-104, Antibodies Inc) and primary rabbit anti-Esr1 antibody (1:1000, 06-935, Millipore) were applied to brain sections from controls, Trpc5^Esr1^-KO and Trpc5^Esr1^-OE mice.

To examine whether CNO increases the activity of Esr1^MPOA^ neurons expressing the AAV-DIO-hM3D(Gq)-mCherry virus, Esr1-Cre mice were injected with either saline or CNO (0.5 mg kg^-1^). Brain sections containing the MPOA were processed for immunofluorescence staining using a primary rabbit anti-c-Fos antibody (1:1000; #2250, Cell Signaling Technology).

To assess whether NachBac induces c-Fos expression in Esr1^MPOA^ neurons, Esr1-Cre male mice were injected with AAV-DIO-NachBac-GFP into the right MPOA. Brain sections containing the MPOA were subsequently processed for immunofluorescence staining using a primary rabbit anti-c-Fos antibody (1:1000; #2250, Cell Signaling Technology).

The following day, sections were rinsed three times for 10 minutes each in 0.1% PBST and then incubated with secondary antibodies: Goat anti-Rabbit 488 (1:500, #111-545-144, Jackson ImmunoResearch), Goat anti-Rabbit Cy3 (1:500, #111-165-144, Jackson ImmunoResearch), Goat anti-Mouse 488 (1:500, #115-545-146, Jackson ImmunoResearch), and Goat anti-Mouse Cy3 (1:500, #115-165-146, Jackson ImmunoResearch) at room temperature for 2 hours with shaking. Sections were cover-slipped and analyzed using a fluorescence microscope. The numbers of each type cell were counted in four or five consecutive brain sections, and the average was used to reflect the data value for each mouse. Three to five mice were included in each group for statistical analyses.

#### RNAscope

Mice were anesthetized and transcardially perfused with 0.9% saline, followed by perfusion with 10% formalin. The brains were then harvested and fixed in 10% formalin at 4°C for 24 hours, followed by cryoprotection in 30% sucrose for 48 hours. The brains were frozen and sectioned into 25 µm slices using a cryostat, and washed in phosphate-buffered saline (PBS) for 10 minutes. Sections were mounted on diethylpyrocarbonate-treated charged slides, dried for 1 h at room temperature and stored at -80 °C. On the day of the RNAscope assay, slides were thawed, rinsed twice in 1× PBS, and placed in a 60°C oven for 30 minutes. Afterward, the slides were post-fixed in 10% formalin at 4°C for 15 minutes. The slides were then dehydrated through graded ethanol (50%, 70%, and 100%, 5 minutes each), followed by target retrieval using microwave treatment at 100°C for 5 minutes. The slides were incubated with Proteinase III (322337, ACDBio) at 40°C for 30 minutes. The slides were then washed with distilled water and incubated with Esr1 (478201-C3, ACDBio) and Trpc5 (503391-C1, ACDBio) RNAscope probes at 40°C for 2.5 hours. Subsequently, the slides were processed using the RNAscope Fluorescent Multiplex Assay Kit (320851, ACDBio) according to the manufacturer’s instructions. Finally, coverslips were applied, and the slides were analyzed using a fluorescence microscope.

#### Fiber photometry

For fiber photometry recordings of Esr1^MPOA^ neurons, eight virgin Esr1-Cre male mice were bilaterally injected with AAV-hSyn-DIO-GCaMP6m. Four of the eight mice exhibited infanticidal behavior four weeks after surgery and were subsequently used for recordings. During the first recording session, animals were left undisturbed in their home cage for approximately 30 minutes before the introduction of a postnatal day 1-3 (P1-3) pup at a location distant from the nest. Following naturally occurring infanticide, the pup was promptly removed and euthanized. The infanticide episode typically lasted 5-10 minutes. To compare Esr1^MPOA^ neuronal activity across different social and non-social stimuli, we sequentially introduced a singly housed adult WT male, a singly housed adult WT female, a novel object, a fake pup, and a water spray into the home cage of the recorded mouse. Each stimulus was presented for 10 minutes with a 5-minute inter-trial interval. After recordings in the hostile virgin state, each male was paired with an adult female until successful mating was confirmed by visible pregnancy and subsequent parturition. On postpartum day 3, the same males (sires) were subjected to recordings using the identical procedures employed during the virgin state. For fiber photometry recordings of deletion Trpc5 in Esr1^MPOA^ neurons, virgin Esr1-Flpo::Trpc5^flox/Y^ male mice paired with an adult female until parturition after the surgery. On PPD3, the sires were subjected to recordings using the identical procedures employed as previously described. For fiber photometry recordings assessing the effect of Trpc5 overexpression in Esr1^MPOA^ neurons, four of the nine control-injected males exhibited infanticidal behavior, whereas four of the seven Trpc5-overexpressing males displayed pup retrieval behavior. These mice were subsequently used for recordings.

Continuous <20 μW blue LED at 465 nm and UV LED at 405 nm served as excitation light sources, driven by a multichannel hub (Doric Lenses), modulated at 211 Hz and 330 Hz, respectively. The light was delivered to a filtered minicube (FMC5, Doric Lenses) before connecting through optic fibers to a rotary joint (FRJ 1 × 1, Doric Lenses) to allow for movement. GCaMP6m or GCaMP6f calcium GFP signals and UV autofluorescent signals were collected through the same fibers back to the dichroic ports of the minicube into a femtowatt silicon photoreceiver (2151, Newport). The digital signals were then amplified, demodulated and collected through a lock-in amplifier (RZ5P, Tucker-Davis Technologies). The fiber photometry data were collected using Doric Synapse v.2.2.1 and sampled down to 8 Hz. We derived the values of calcium fluorescence change (ΔF/F) by calculating (F_465_-F_0_)/F_0_, where F_0_ is the baseline fluorescence of the F_465_ channel signal 4 s before and after. A 405 nm signal as a control for 465 nm.

#### Slice electrophysiology

Electrophysiological recordings were performed using a protocol described previously^23^. Electrophysiological recordings were performed in hostile WT virgin males, WT sires, Trpc5^Esr1^-KO mice, Trpc5^Esr1^-OE mice, and Esr1-Cre mice injected with hM3D(Gq)-mCherry. Mice were deeply anesthetized with isoflurane and transcardially perfused with a modified ice-cold sucrose-based cutting solution (pH 7.3) containing 10 mM NaCl, 25 mM NaHCO3, 195 mM Sucrose, 5 mM Glucose, 2.5 mM KCl, 1.25 mM NaH2PO4, 2 mM Na-Pyruvate, 0.5 mM CaCl2, and 7 mM MgCl2, bubbled continuously with 95% O2 and 5% CO2. The mice were then decapitated, and the entire brain was removed and immediately submerged in the cutting solution. Slices (220 µm) were cut with a Microm HM 650V vibratome (Thermo Scientific). Two brain slices containing the MPOA were obtained for each animal (bregma 0.14 mm to −0.22 mm). The slices were recovered for 30 min at 34°C and then maintained at room temperature in artificial cerebrospinal fluid (aCSF, pH 7.3) containing 126 mM NaCl, 2.5 mM KCl, 2.4 mM CaCl2, 1.2 mM NaH2PO4, 1.2 mM MgCl2, 11.1 mM glucose, and 21.4 mM NaHCO3) saturated with 95% O2 and 5% CO2 before recording. Slices were transferred to a recording chamber and allowed to equilibrate for at least 10 min before recording. The slices were superfused at 34°C in oxygenated aCSF at a flow rate of 1.8-2 ml/min. GFP labeled neurons in the MPOA were visualized using epifluorescence and infrared–differential interference contrast (IR-DIC) imaging on an upright microscope (Eclipse FN-1, Nikon) equipped with a moveable stage (MP-285, Sutter Instrument). Patch pipettes with resistances of 3–5 MΩ were filled with intracellular solution (pH 7.3) containing 128 mM potassium gluconate, 10 mM KCl, 10 mM HEPES, 0.1 mM EGTA, 2 mM MgCl2, 0.05 mM (Na)2GTP, and 0.05 mM (Mg)ATP. Recordings were made using a MultiClamp 700B amplifier (Axon Instruments), sampled using Digidata 1440A, and analyzed offline with pClamp 10.3 software (Axon Instruments). Series resistance was monitored during the recording, and the values were generally <10 MΩ and were not compensated. The liquid junction potential was +12.5 mV and was corrected after the experiment. Data were excluded if the series resistance exceeding 20% changed during the experiment or without overshoot for action potential. Currents were amplified, filtered at 1 kHz, and digitized at 10 kHz. The current-clamp mode was engaged to test neural firing frequency and resting membrane potential (RM). To determine the intrinsic excitability of Esr1^MPOA^ cells, we performed current-clamp recordings and injected 30 current steps ranging from -20 pA to 270 pA in 10 pA increments into the recorded cell. The total number of spikes during each 500-ms-long current step was then used to construct the F-I curve. To test CNO activation of Esr1^MPOA^ neurons, current-clamp recordings measured firing frequency and resting membrane potential (RMP) at baseline and after CNO application (10 μM, 5 s puff). CNO was delivered using a consistent pressure system (Picospritzer III). Neurons with stable 1-minute baselines were included. RMP and firing rates were averaged over the baseline and a 1-minute window centered on the maximal post-CNO response. To test NachBac-induced activation of Esr1^MPOA^ neurons, we performed current-clamp recordings to measure firing frequency and RMP at baseline in GFP-labeled neurons from Esr1-Cre mice that received either AAV-DIO-GFP or AAV-DIO-NachBac-GFP injection in the MPOA.

### Statistical analysis

All data are presented as the means ± SEM. Statistical analysis was performed using GraphPad Prism 10.0 (GraphPad Software, Inc). Two-tailed unpaired t-test, paired t-test, chi-squared test and Mann-Whitney test (if data were not normally distributed) were used to determine significant differences between the two groups. Two-way ANOVA ± Sidak corrections were used for comparisons involving two variables ± multiple testing (e.g. genotype and time). The difference between the three or above groups was tested using the one-way ANOVA. A significance (alpha) level of p < 0.05 was considered statistically significant.

## Acknowledgement

The investigators were supported by grants from the NIH (R01HD114146 to YX), American Diabetes Association (1-24-PDF-56 to YL), and the Silver Endowment (to YX). ISF is supported by Wellcome (207462/Z/17/Z), Fondation Botnar, the Bernard Wolfe Health Neuroscience Endowment, the Leducq Foundation and a NIHR Senior Investigator Award.

## Author contributions

YL and YX conceived the project, experimental design and wrote the manuscript. YL performed the procedures, data acquisition and analyses. QL performed electrophysiology studies and analyzed the data. FW, KMM, MW, YD, Yuxue Yang, Yutian Liu, JC, MS, XL, JJ, JQ, XW, LX and TZ contributed to the generation of study mice and data discussion. Yongjie Yang, HL, LT, BRA and ISF were involved in study design and data discussion.

## Data and materials availability

All data used in the analysis is available to any researcher for purposes of reproducing or extending the analysis.

---

**Extended data Figure 1.**
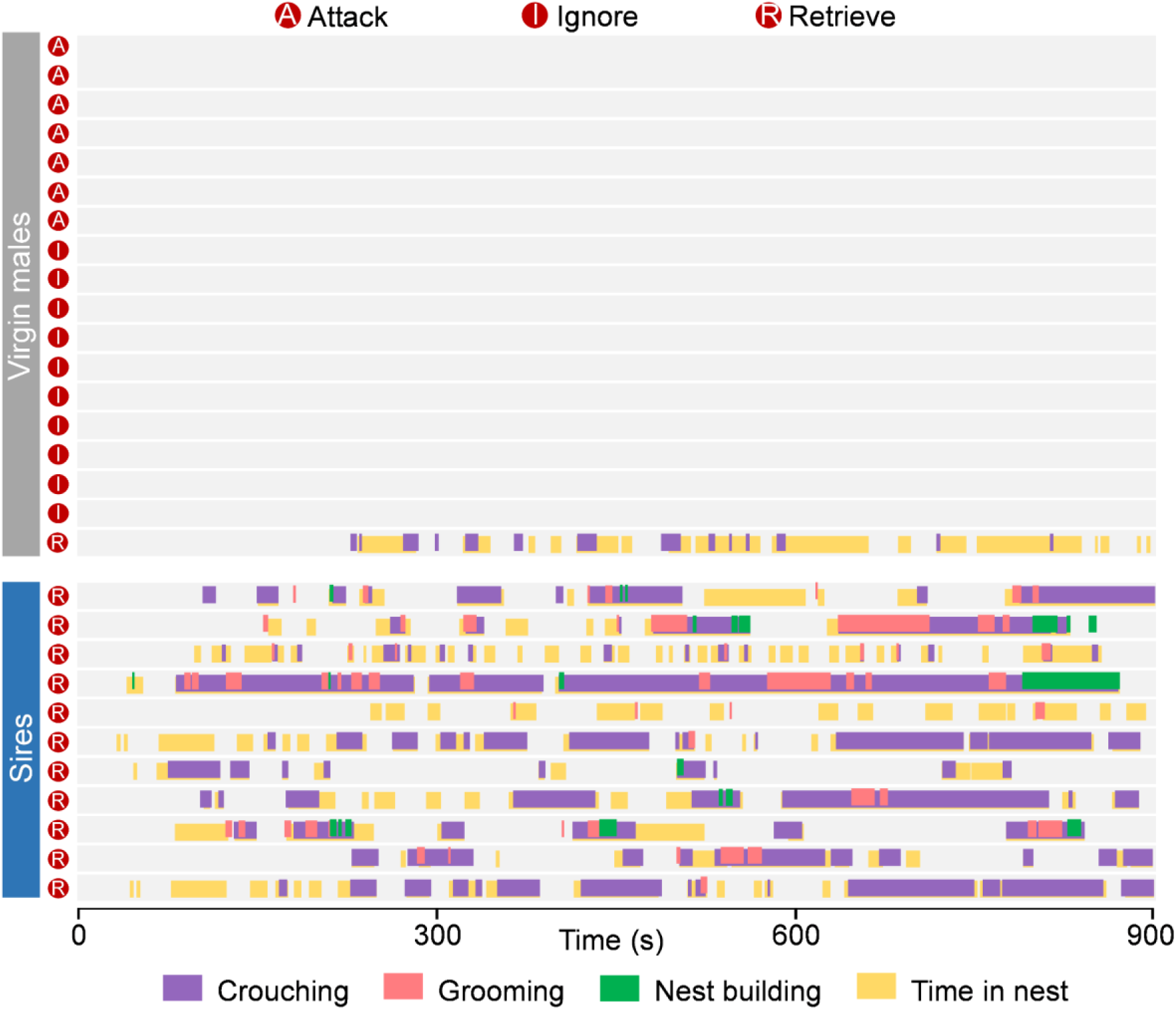
Behavior raster plot of WT virgin males and sires that attack, ignore or retrieve pups.

**Extended data Figure 2.**
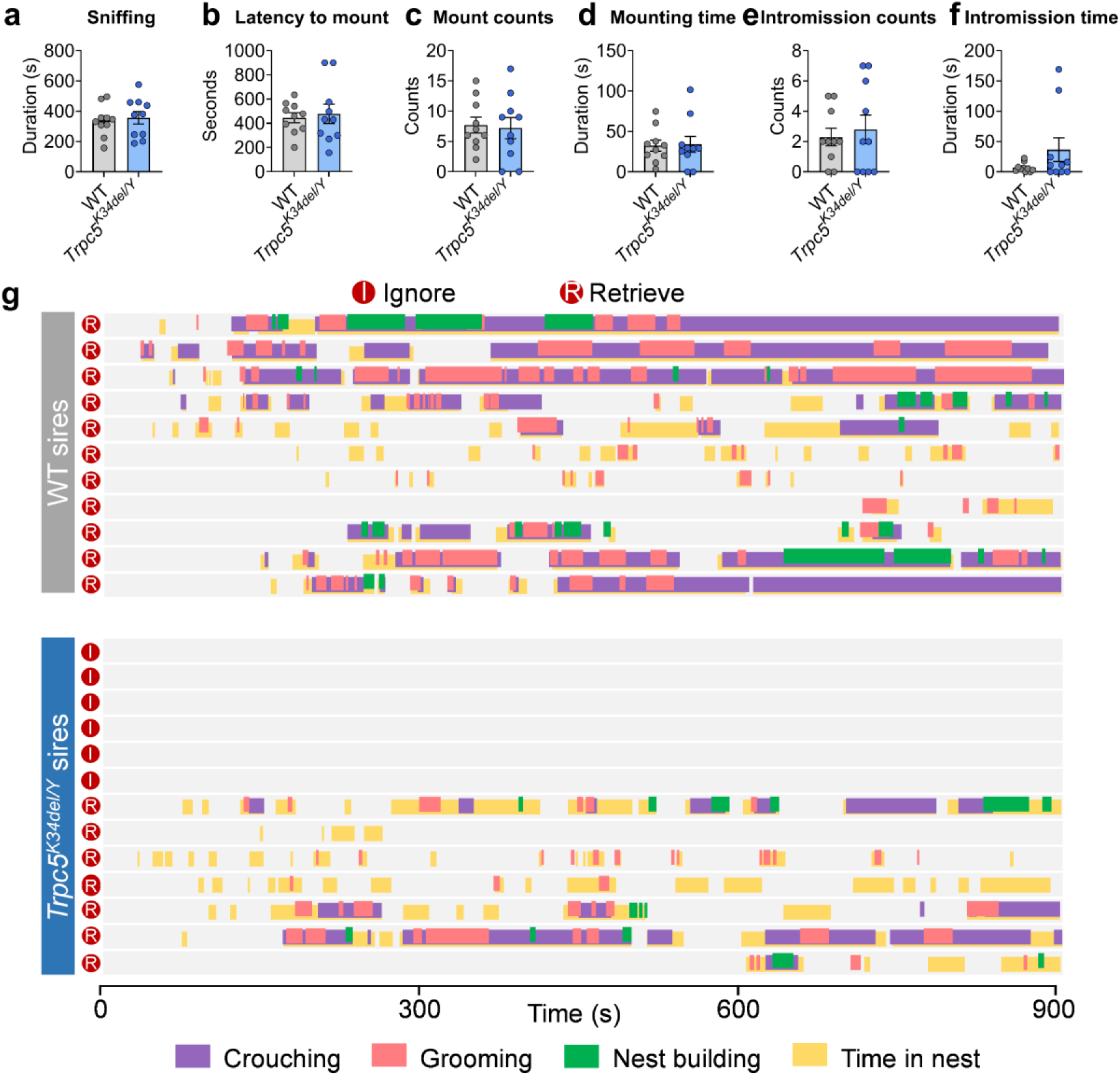
Sexual behaviors of WT and Trpc5^K34del/Y^ mice and behavior raster plot of sires. (a-f) Sexual behavior of WT and Trpc5^K34del/Y^ virgin male mice. Sniffing the female mice (a), latency to mount the female mice (b), mount counts (c), mounting time (d), intromission counts (e) and intromission time (f). (g) Behavior raster plot of WT and Trpc5^K34del/Y^ mice that show ignore or retrieve. Data presented as mean ± SEM. p value determined using unpaired t test (a,b) and Mann-Whitney test (c,d,e,f).

**Extended data Figure 3.**
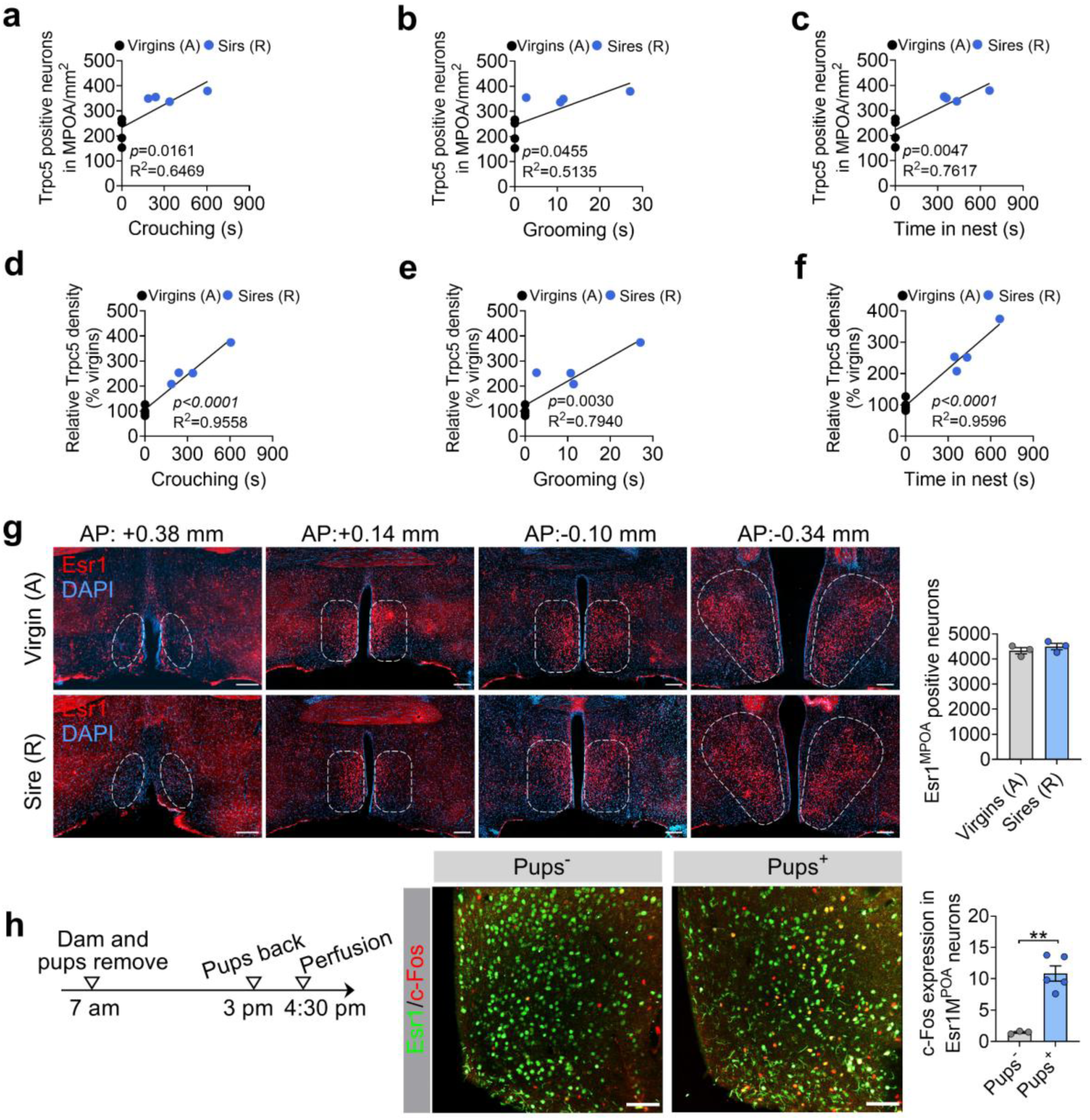
Correlation between Trpc5 expression and paternal behavior parameters, characterization of Esr1^MPOA^ neurons in hostile virgin males and sires, and pup-induced activation of Esr1^MPOA^ neurons. (a-c) Positive correlations between the number of Trpc5+ neurons in the MPOA and three paternal behavior parameters: crouching time (a), grooming time (b), and time spent in the nest (c) in hostile virgin males and pup-retrieving sires. Each dot represents one animal (n = 4 per group, 12 weeks of age). (d-f) Similar correlations were observed between the density of Trpc5 immunoreactivity in the MPOA and crouching (d), grooming (e), and nest time (f). Each dot represents one animal (n = 4 per group, 12 weeks of age). (g) Left: representative images showing Esr1 neurons (red) across anterior-posterior levels of the MPOA in hostile virgin males (top) and pup-retrieving sires (bottom). Right: quantification shows no significant difference in the total number of Esr1^MPOA^ neurons between groups (n = 4 per group). Scale bars, 100 μm. (h) Pup-induced c-Fos expression in Esr1^MPOA^ neurons. Experimental timeline: dams and pups were removed at 7:00 a.m., and pups were returned at 3:00 p.m.; mice were perfused at 4:30 p.m. Representative images show Esr1 (green) and c-Fos (red) co-expression in MPOA of males with or without pup exposure. Quantification reveals a significant increase in c-Fos-Esr1^MPOA^ neurons in the pup-exposed group (n = 4 per group, 18 weeks of age). Scale bars, 50 μm. Data presented as mean ± SEM. p value determined using unpaired t-test (g,h). **p < 0.01

**Extended data Figure 4.**
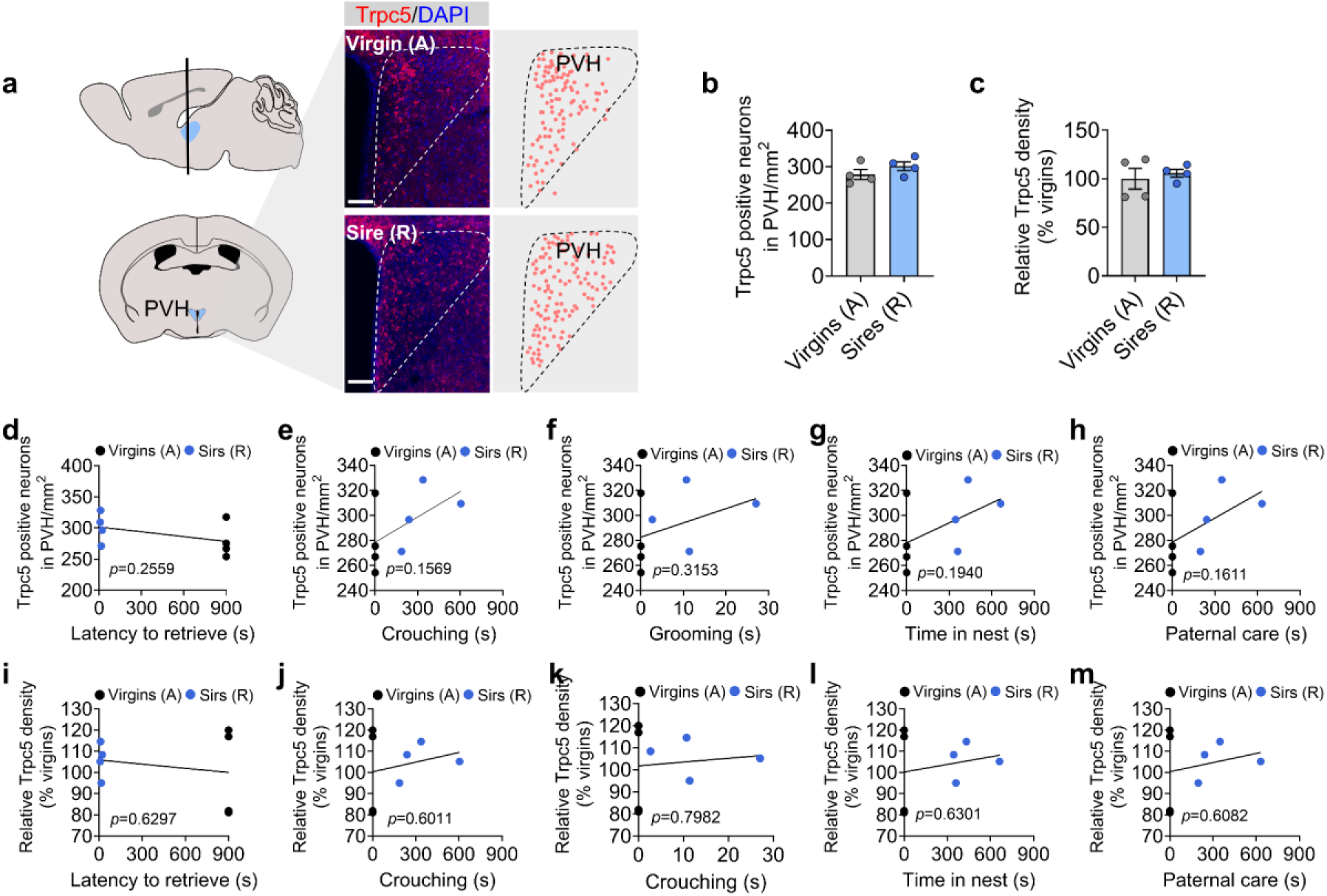
Fatherhood experience did not change Trpc5 expression in the PVH. (a) Immunofluorescence images showing Trpc5 expression in the PVH of hostile virgin males and pup-retrieving sires (n = 4 per group, 12 weeks of age). Scale bar, 60 μm. (b-c) Quantification of Trpc5+ neuron number (b) and density of Trpc5 immunoreactivity (c) in the PVH of hostile virgin males and pup-retrieving sires (n = 4 per group, 12 weeks of age). (d-h) Correlation between the number of Trpc5+ neurons in the PVH and behavioral performance, including latency to retrieve pups (d), crouching (e), grooming (f), time in nest (g) and paternal care duration (h). (i-m) Correlation between the density of Trpc5 immunoreactivity in the PVH and behavioral performance, including latency to retrieve pups (i), crouching (j), grooming (k), time in nest (l) and paternal care duration (m). Each dot represents one animal (n = 4 per group, 12 weeks of age). Data presented as mean ± SEM. p value determined using unpaired t test (b,c).

**Extended data Figure 5.**
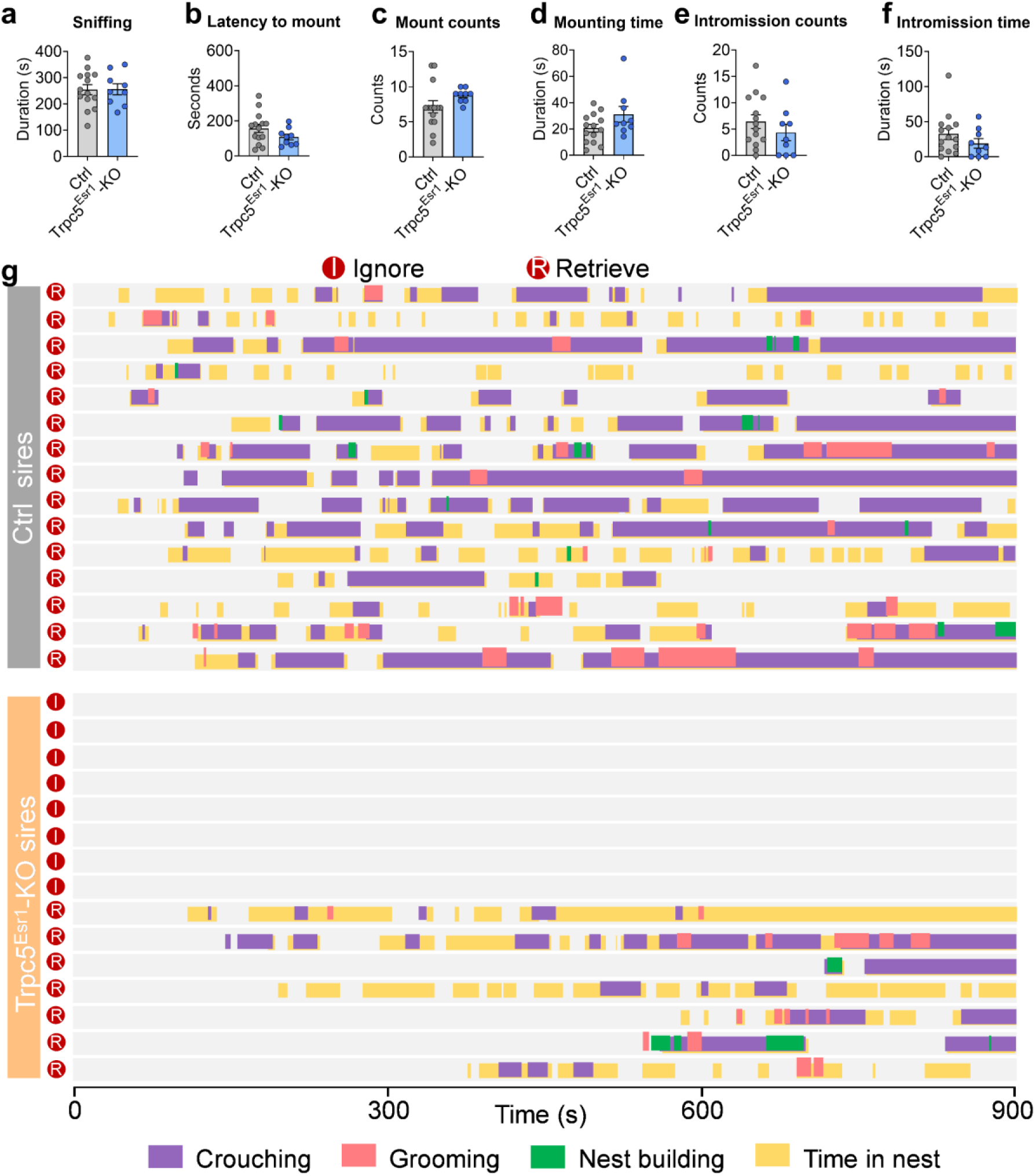
Sexual behaviors of control and Trpc5^Esr1^-KO mice and behavior raster plot of sires. (a-f) Sexual behavior of control and Trpc5^Esr1^-KO mice virgin male mice. Sniffing the female mice (a), latency to mount the female mice (b), mount counts (c), mounting time (d), intromission counts (e) and intromission time (f). (g) Behavior raster plot of control and Trpc5^Esr1^-KO sires that show ignore or retrieve. Data presented as mean ± SEM. p value determined using unpaired t-test (a,b) and Mann-Whitney test (c,d,e,f).

**Extended data Figure 6.**
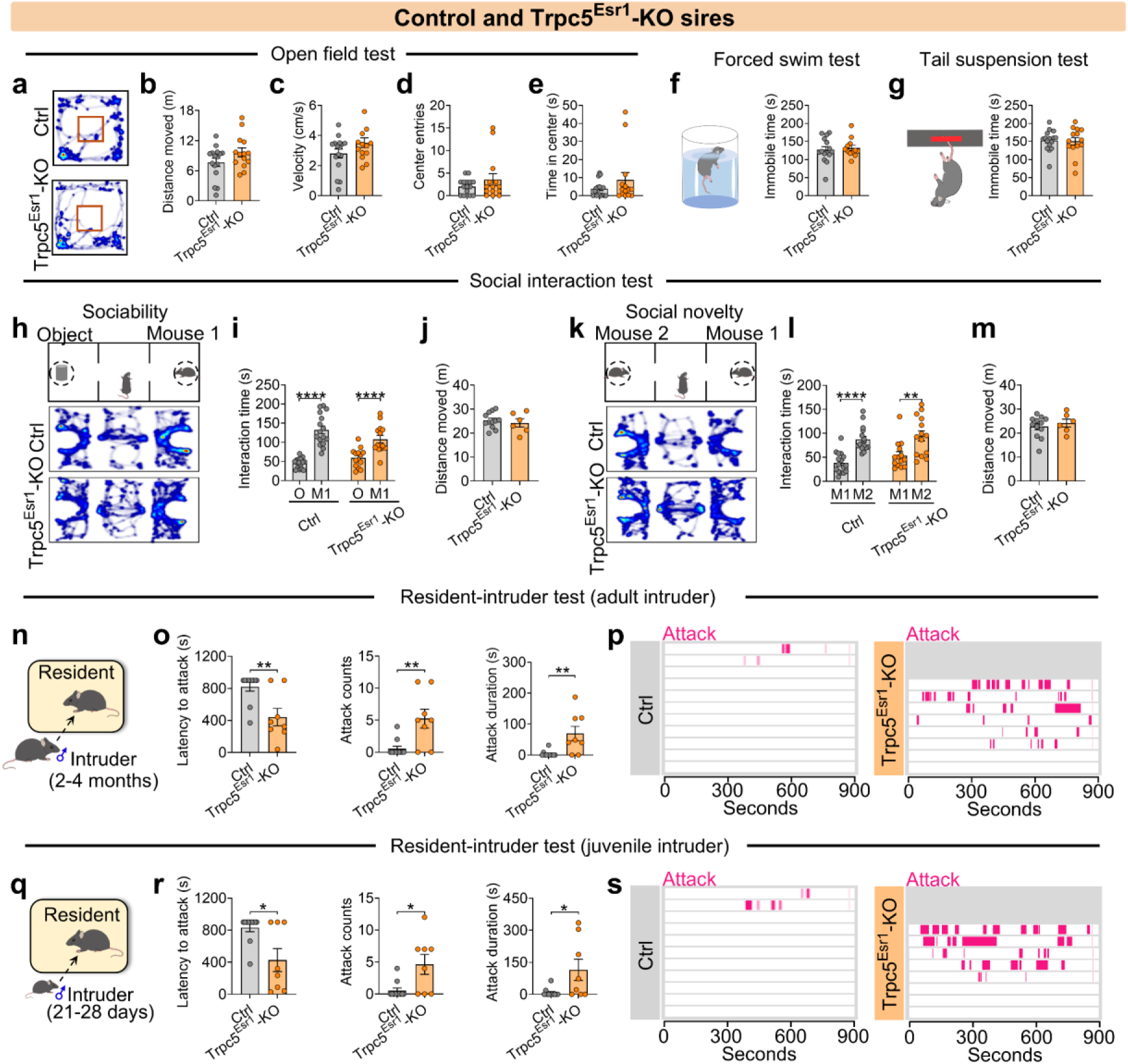
Other social behaviors of control and Trpc5^Esr1^-KO sires. (a-e) Heatmap of movement in the open field test (a), total distance moved (b), velocity (c), center entries (d) and center duration (e) between control and Trpc5^Esr1^-KO sires (n=14-16 mice per group, 18 weeks of age). (f-g) Immobile time in forced swim test (f) and tail suspension test (g) between control and Trpc5^Esr1^-KO sires (n=12-15 mice per group, 18 weeks of age). (h) Schematic diagram of three-chamber social ability test. (i) Interaction time with object and mouse 1 in chamber by control and Trpc5^Esr1^-KO sires in social ability test (n = 14-16 mice per group, 19 weeks of age). (j) Distances moved in three chambers by control and Trpc5^Esr1^-KO sires in social ability test (n = 6-12 mice per group, 19 weeks of age). (k) Schematic diagram of three-chamber social novelty test. (l) Interaction time with mouse 2 and mouse a in chamber by control and Trpc5^Esr1^-KO sires in social novelty test (n = 14-16 mice per group, 19 weeks of age). (m) Distances moved in three chambers by control and Trpc5^Esr1^-KO sires in social novelty test (n = 6-12 mice per group, 19 weeks of age). (n) Schematic diagram of resident-intruder assay (Group housed adult male intruder mice). (o) Latency to attack (left), number of attacks (middle), and duration of attack (right) between control and Trpc5^Esr1^-KO sires (n = 8-11 mice per group, 20 weeks of age). (p) Raster plots showing inter-male aggressive behavior. (q) Schematic diagram of resident-intruder assay (Group housed juvenile male intruder mice). (r) Latency to attack (left), number of attacks (middle), and duration of attack (right) between control and Trpc5^Esr1^-KO sires (n = 8-11 mice per group, 20 weeks of age). (s) Raster plots showing inter-male aggressive behavior. Data presented as mean ± SEM, p value determined using unpaired t-tests (b,c,d,e,f,g,i,j,l,m) and Mann-Whitney test (o,r). *p < 0.05, **p < 0.01 and ****p < 0.0001.

**Extended data Figure 7.**
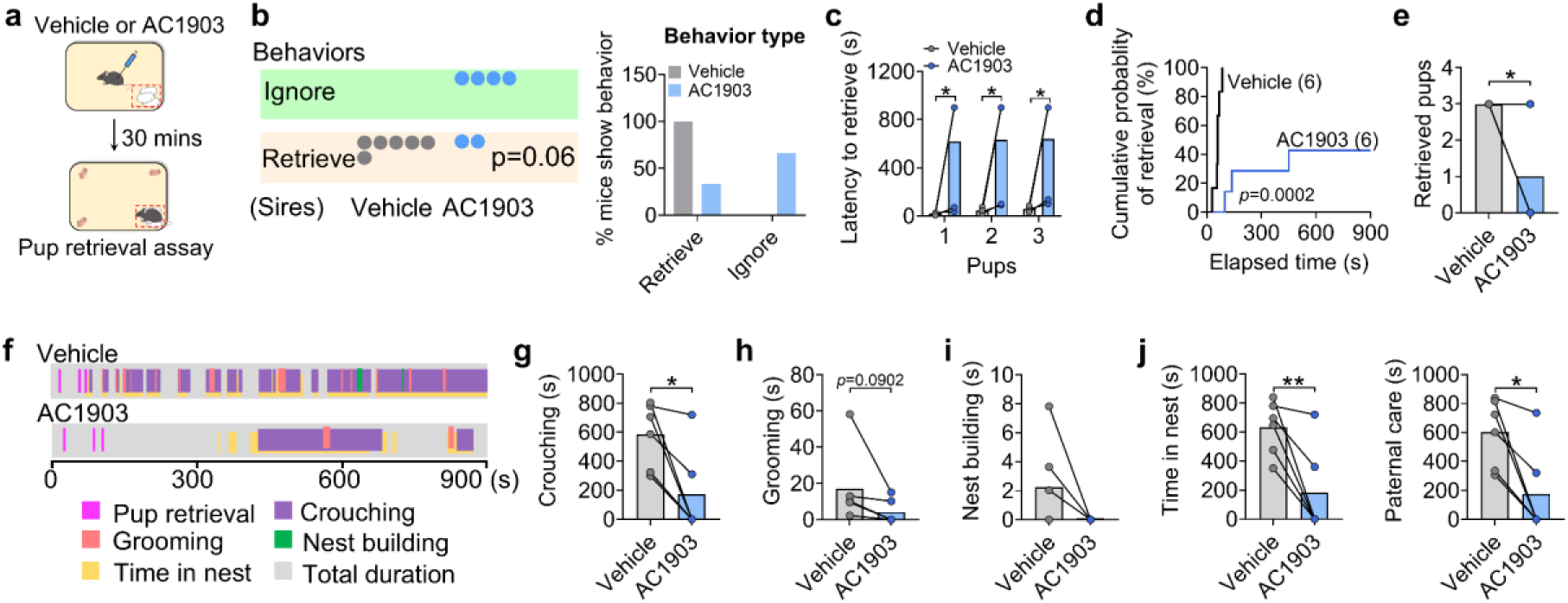
Pharmacological inhibition of Trpc5 impairs paternal behavior in WT sires. (a) The paternal behavior of WT sires receiving vehicle or AC1903 (the Trpc5 inhibitor) injection were studied in the home cage. (b) Dots indicate the number of WT sires receiving vehicle or AC1903 that exhibited ignoring or retrieving (n= 6 mice per group, 20 weeks of age). (c) The latency to retrieve pups. (d) Cumulative probability of pup retrieval. (E) The numbers of retrieved pups. (f) Representative behavior raster plots of two individual mouse (vehicle and AC1903) in the pup-directed behavior. (g-k) The duration of crouching pups (g), grooming pups (h), nest building (i), time spent in the nest (j) and paternal care duration (k) (n = 6 mice per group, 20 weeks of age). Data presented as mean ± SEM. p value determined using paired t-test (c,e,g,h,i,j,k) or chi-squared test (b). *p < 0.05 and **p < 0.01.

**Extended data Figure 8.**
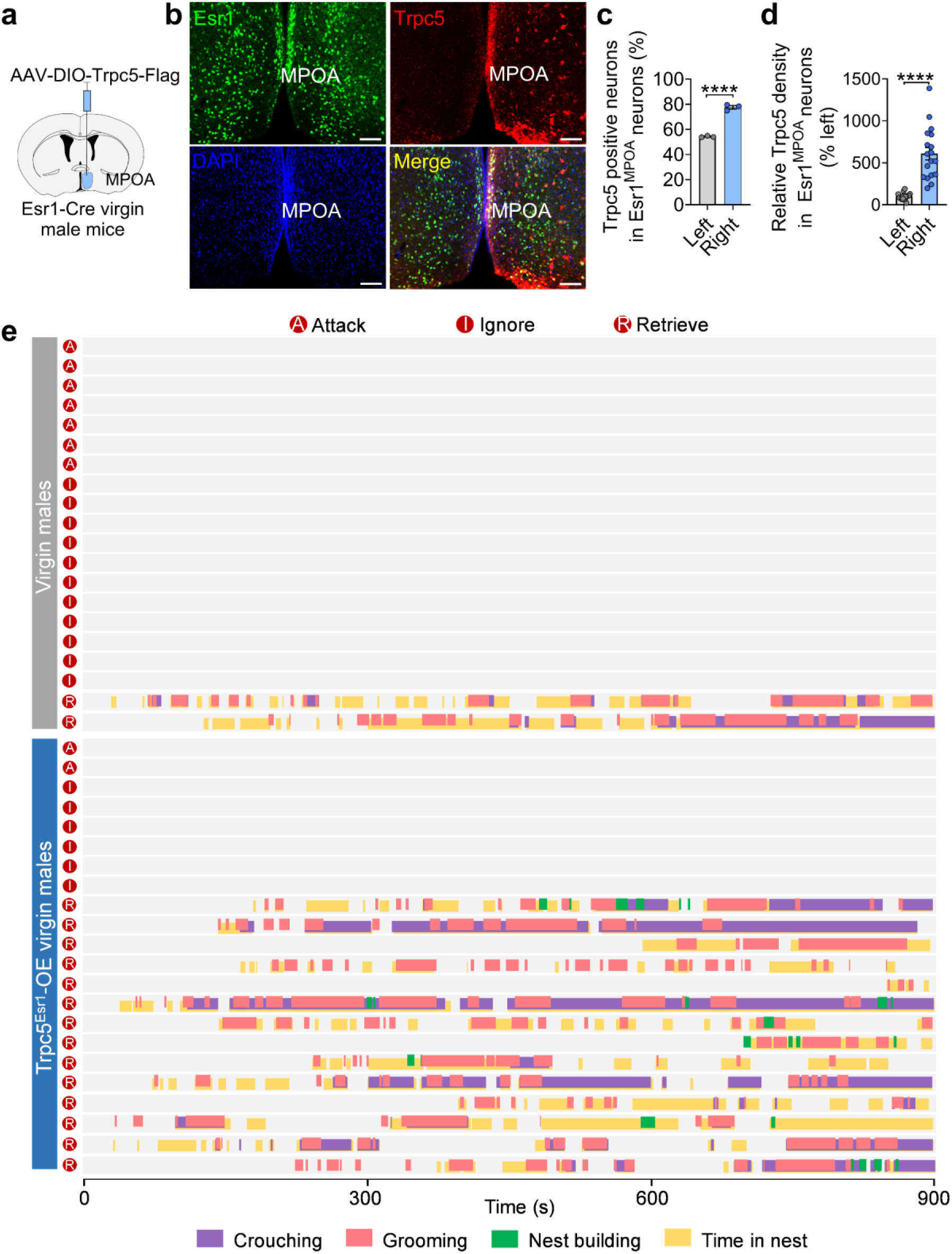
Validation of Trpc5 overexpression and behavior raster plot of control and Trpc5^Esr1^-OE virgin male mice that attack, ignore or retrieve. (a) Schematic diagram of validation of Trpc5 overexpression using AAV-DIO-Trpc5-Flag (injection of right side of MPOA). (b) Representative images showing the expression of Trpc5 in Esr1^MPOA^ neurons after injection of AAV-DIO-Trpc5-Flag. (c) The percentage of Esr1^MPOA^ neurons (green) co-expressing Trpc5 (red) in Esr1-Cre virgin male mice receiving AAV-DIO-Trpc5 in the right MPOA (*n* = 3 mice, 15 weeks of age). Scale bars, 100 μm. Nuclei are counterstained with DAPI (blue). (d) Quantification of Trpc5 density within Esr1^MPOA^ neurons. (e) Behavior raster plot of control and Trpc5^Esr1^-OE virgin male mice that attack, ignore or retrieve. Data presented as mean ± SEM. p value determined using unpaired t-test (c,d). ****p < 0.0001.

**Extended data Figure 9.**
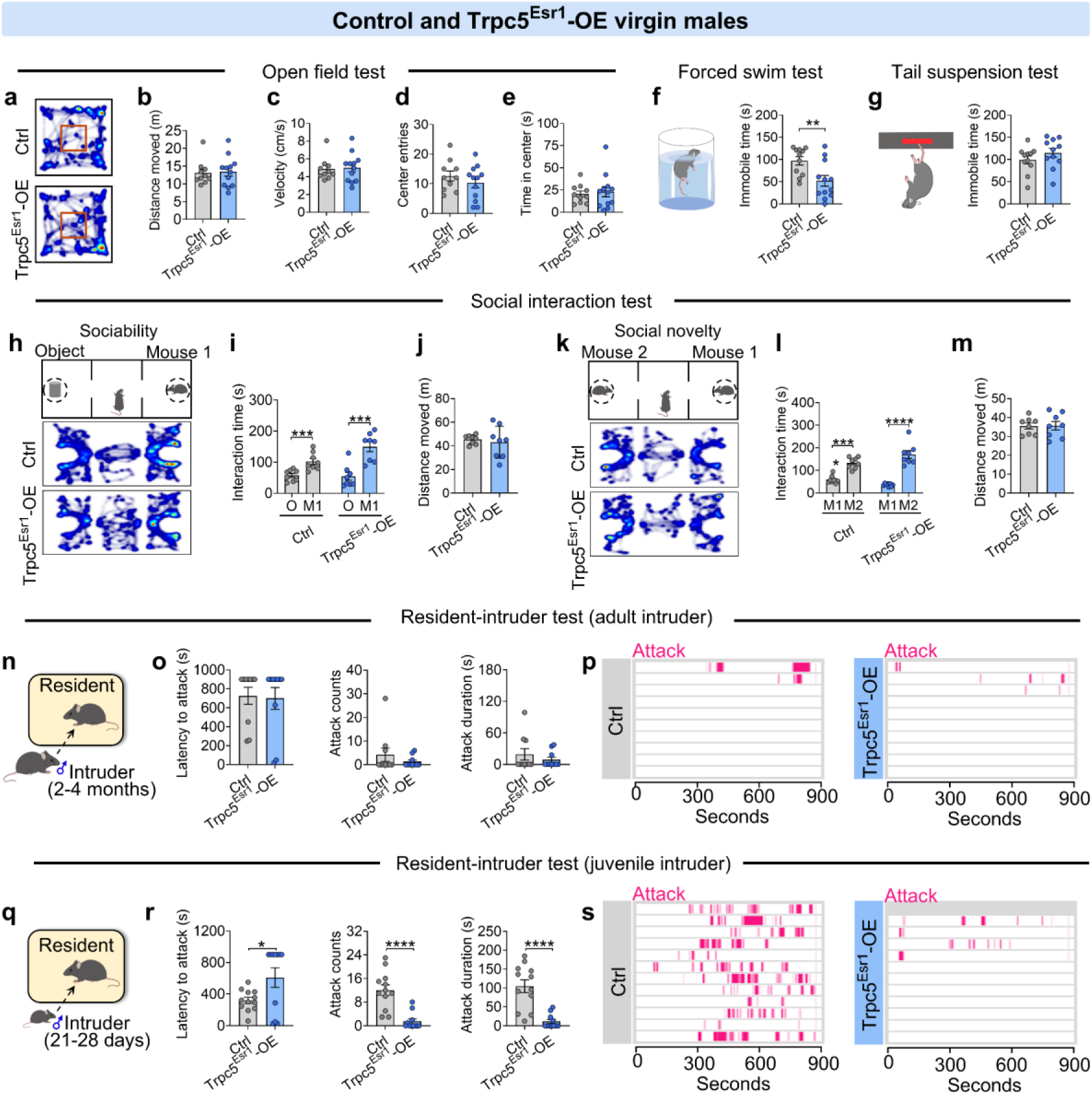
Other social behaviors of control and Trpc5^Esr1^-OE virgin males. (a-e) Heatmap of movement in the open field test (a), total distance moved (b), velocity (c), center entries (d) and center duration (e) between control and Trpc5^Esr1^-OE virgin male mice (n=10-12 mice per group, 15 weeks of age). (f-g) Immobile time in forced swim test (f) and tail suspension test (g) between control and Trpc5^Esr1^-OE virgin male mice (n=11-12 mice per group, 16 weeks of age). (h) Schematic diagram of three-chamber social ability test. (i) Interaction time with object and mouse 1 in chamber control and Trpc5^Esr1^-OE virgin male mice in social ability test (n = 8 mice per group, 17 weeks of age). (j) Distances moved in three chambers by control and Trpc5^Esr1^-KO father mice in social ability test (n = 8 mice per group, 17 weeks of age). (k) Schematic diagram of three-chamber social novelty test. (l) Interaction time with mouse 2 and mouse a in chamber control and Trpc5^Esr1^-OE virgin male mice in social novelty test (n = 8 mice per group, 17 weeks of age). (m) Distances moved in three chambers control and Trpc5^Esr1^-OE virgin male mice in social novelty test (n = 8 mice per group, 17 weeks of age). (n) Schematic diagram of resident-intruder assay (Group housed adult male intruder mice). (o) Latency to attack (left), number of attacks (middle), and duration of attack (right) between control and Trpc5^Esr1^-OE virgin male mice (n = 10 mice per group, 18 weeks of age). (p) Raster plots showing inter-male aggressive behavior. (q) Schematic diagram of resident-intruder assay (Group housed juvenile male intruder mice). (r) Latency to attack (left), number of attacks (middle), and duration of attack (right) between control and Trpc5^Esr1^-OE virgin male mice (n = 11-12 mice per group, 18 weeks of age). (s) Raster plots showing inter-male aggressive behavior. Data presented as mean ± SEM, p value determined using unpaired t-tests (b,c,d,e,f,g,j,m) and Mann-Whitney test (o,r). *p < 0.05, ***p < 0.001 and ****p < 0.0001.

**Extended data Figure 10.**
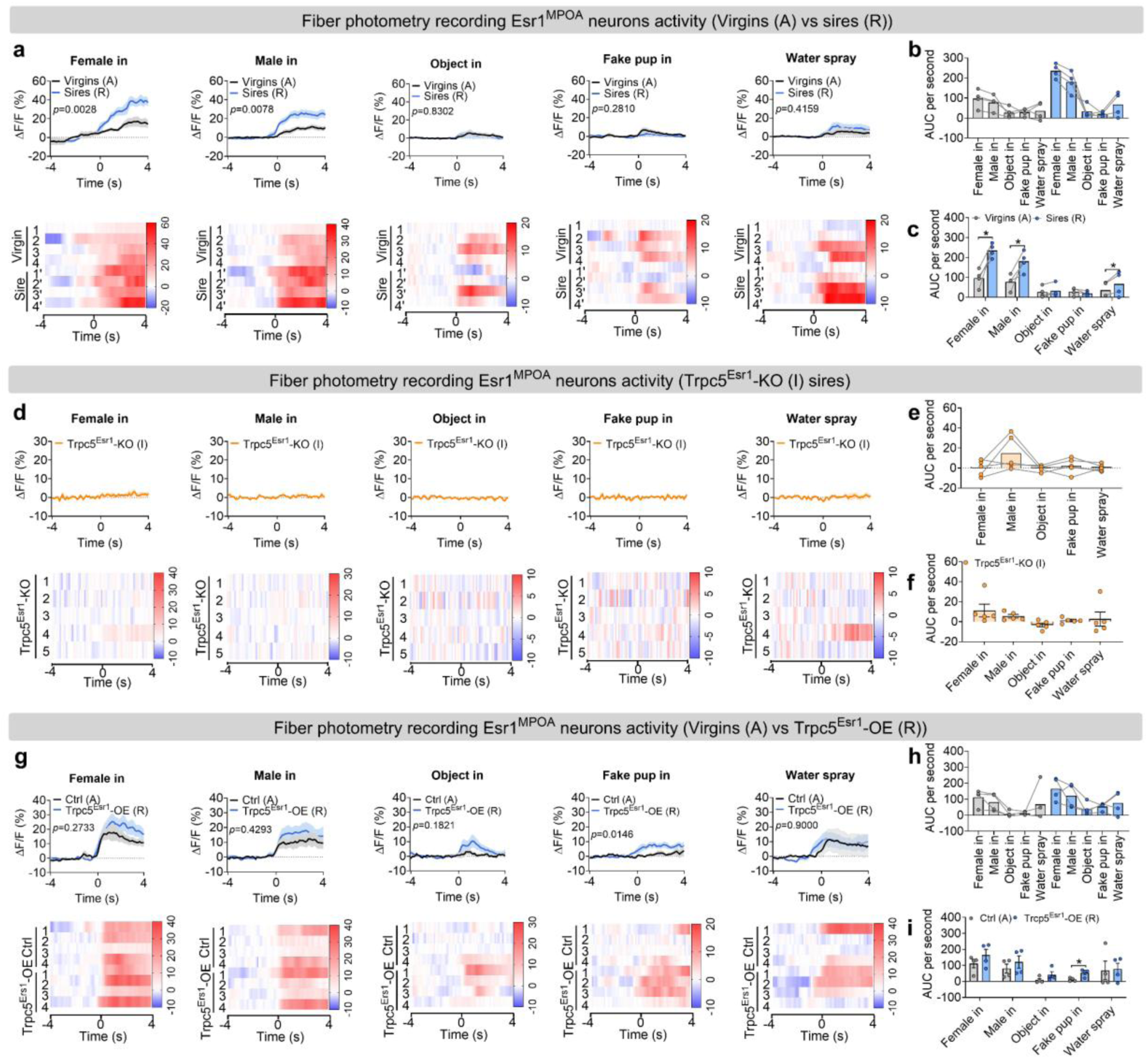
Esr1^MPOA^ neuronal responses during the transition to fatherhood and the effects of Trpc5 manipulation to other social and non-social cues. (a) The ΔF/F calcium signal (top) and heatmap (bottom) of Esr1^MPOA^ neurons aligned to the onset of the following behaviors: female intruder in, male intruder in, object in, fake pup in and water spray (from left to right). (b-c) The mean AUC of the ΔF/F during various social and non-social behaviors to compare responses across behaviors in hostile virgin males and sires (b) and responses of the same behavior between hostile virgin males and sires (c) (n = 4 mice, 12 weeks of age). (d) The ΔF/F calcium signal (top) and heatmap (bottom) of Esr1^MPOA^ neurons aligned to the onset of the following behaviors: female intruder in, male intruder in, object in, fake pup in and water spray (from left to right). (e-f) The mean AUC of the ΔF/F during various social and non-social behaviors to compare responses across behaviors in Trpc5^Esr1^-KO sires (e) and responses of the same behavior in Trpc5^Esr1^-KO sires (f) (n = 5 mice, 18 weeks of age). (g) The ΔF/F calcium signal (top) and heatmap (bottom) of Esr1^MPOA^ neurons aligned to the onset of the following behaviors: female intruder in, male intruder in, object in, fake pup in and water spray (from left to right). (h-i) The mean AUC of the ΔF/F during various social and non-social behaviors to compare responses across behaviors in hostile virgin males and Trpc5^Esr1^-OE virgin males (h) and responses of the same behavior in hostile virgin males and Trpc5^Esr1^-OE virgin males (i) (n = 4 mice, 12 weeks of age). Data presented as mean ± SEM, p value determined using two-way ANOVA (b,e,h), paired t-test (c), unpaired t-test (f,i). *p < 0.05.

**Extended data Figure 11.**
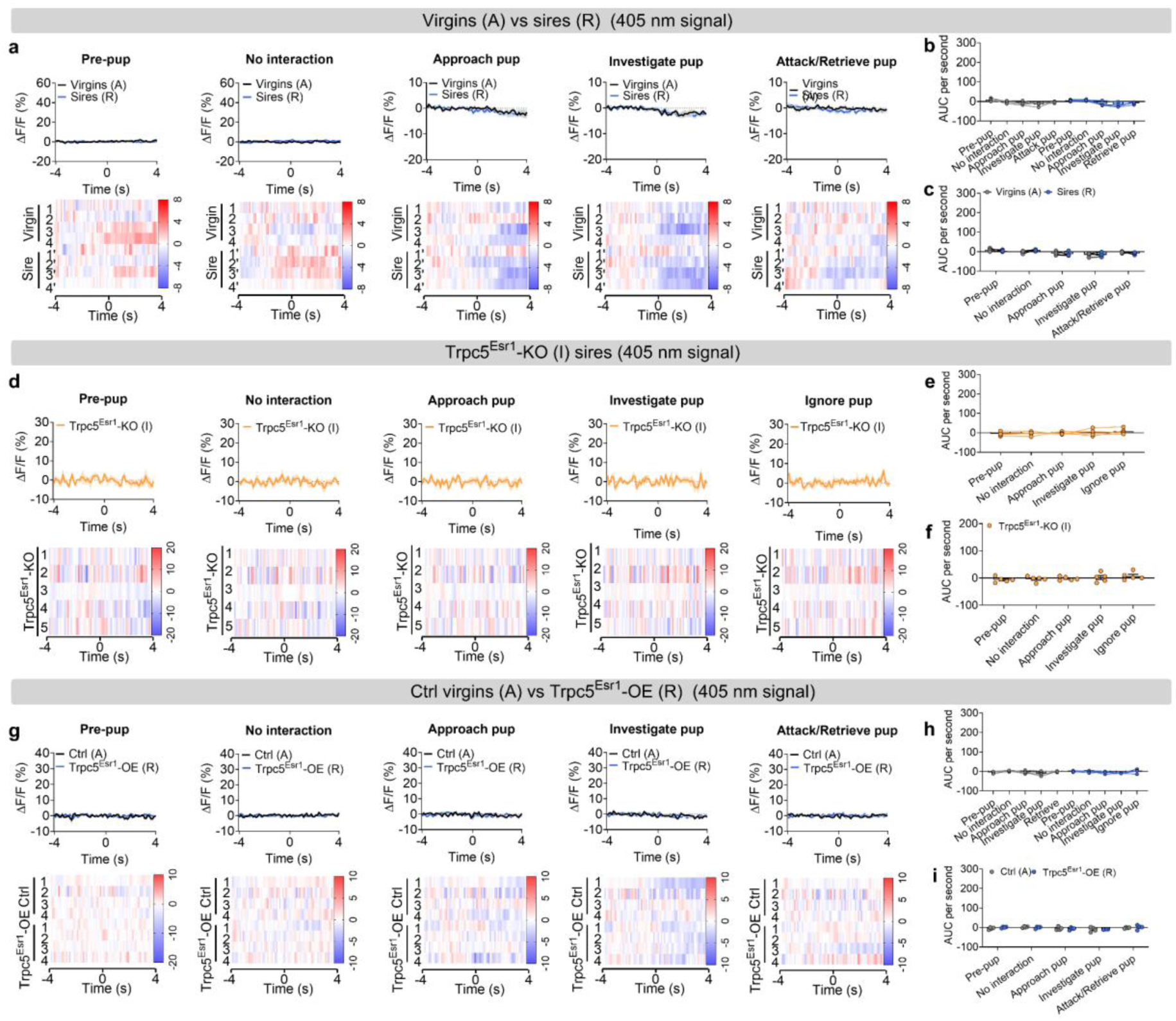
The 405 nm control signal from Esr1^MPOA^ neuronal recordings unchanged in response to pup cues. (a) The 405 nm control ΔF/F calcium signal (top) and heatmap (bottom) of Esr1^MPOA^ neurons aligned to the onset of the following behaviors: pre-pup, no interaction, approach pup, investigate pup and attack/retrieve pup (from left to right). (b-c) The mean AUC of the ΔF/F during various behaviors to compare responses across behaviors in hostile virgin males and sires (b) and responses of the same behavior between hostile virgin males and sires (c) (n = 4 mice, 12 weeks of age). (d) The 405 nm control ΔF/F calcium signal (top) and heatmap (bottom) of Esr1^MPOA^ neurons aligned to the onset of the following behaviors: pre-pup, no interaction, approach pup, investigate pup and ignore pup (from left to right). (e-f) The mean AUC of the ΔF/F during various behaviors to compare responses across behaviors in Trpc5^Esr1^-KO sires (e) and responses of the same behavior in Trpc5^Esr1^-KO sires (f) (n = 5 mice, 18 weeks of age). (g) The 405 nm control ΔF/F calcium signal (top) and heatmap (bottom) of Esr1^MPOA^ neurons aligned to the onset of the following behaviors: pre-pup, no interaction, approach pup, investigate pup and attack/retrieve pup (from left to right). (h-i) The mean AUC of the ΔF/F during various behaviors to compare responses across behaviors in hostile virgin males and Trpc5^Esr1^-OE virgin males (h) and responses of the same behavior in hostile virgin males and Trpc5^Esr1^-OE virgin males (i) (n = 4 mice, 12 weeks of age). Data presented as mean ± SEM, p value determined using two-way ANOVA (b,e,h), paired t-test (c), unpaired t-test (f,i).

**Extended data Figure 12.**
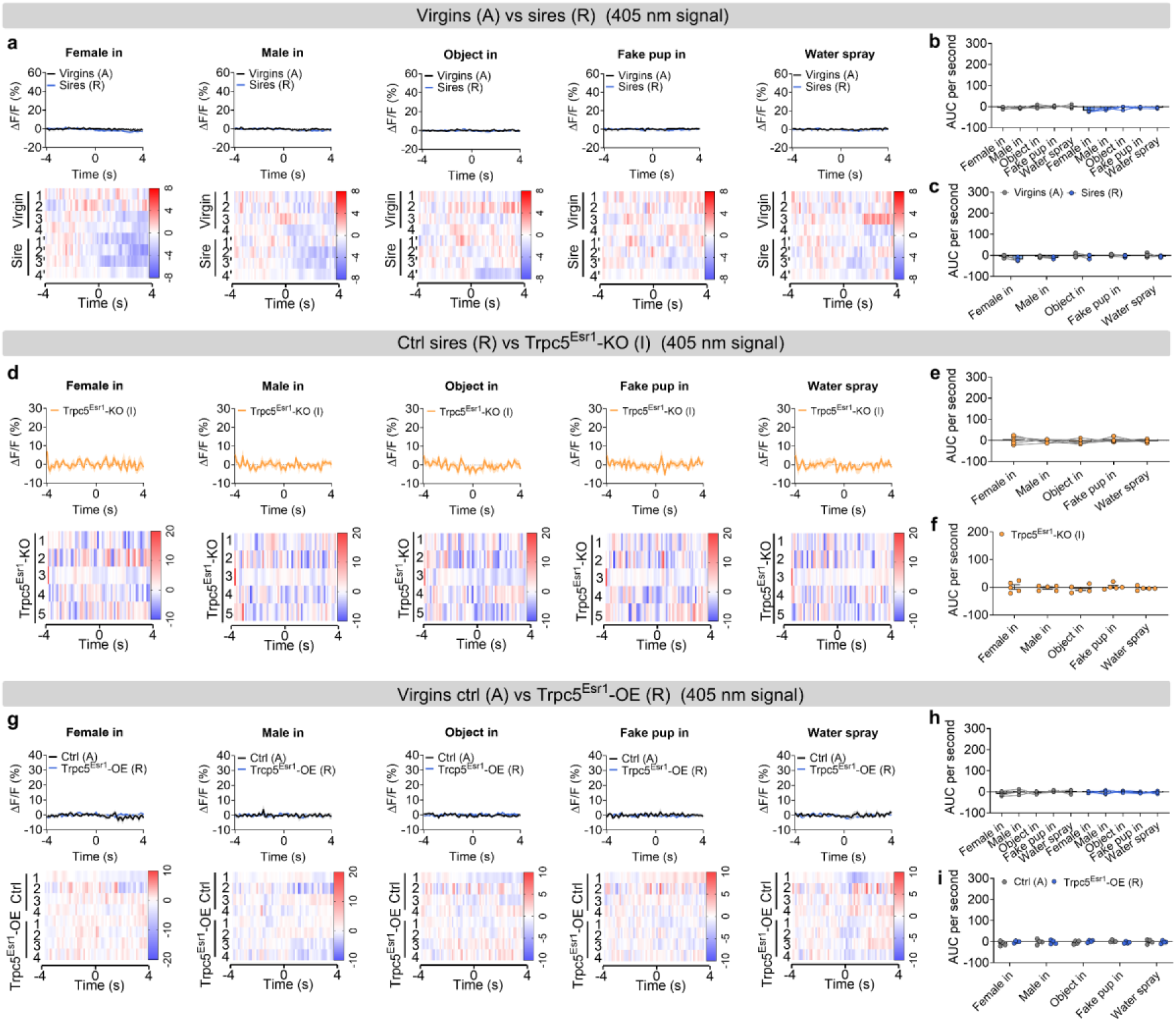
The 405 nm control signal from Esr1^MPOA^ neuronal recordings unchanged in response to other social and non-social cues. (a) The 405 nm control ΔF/F calcium signal (top) and heatmap (bottom) of Esr1^MPOA^ neurons aligned to the onset of the following behaviors: female intruder in, male intruder in, object in, fake pup in and water spray (from left to right). (b-c) The mean AUC of the ΔF/F during various social and non-social behaviors to compare responses across behaviors in hostile virgin males and sires (b) and responses of the same behavior between hostile virgin males and sires (c) (n = 4 mice, 12 weeks of age). (d) The 405 nm control ΔF/F calcium signal (top) and heatmap (bottom) of Esr1^MPOA^ neurons aligned to the onset of the following behaviors: female intruder in, male intruder in, object in, fake pup in and water spray (from left to right). (e-f) The mean AUC of the ΔF/F during various social and non-social behaviors to compare responses across behaviors in Trpc5^Esr1^-KO sires (e) and responses of the same behavior in Trpc5^Esr1^-KO sires (f) (n = 5 mice, 18 weeks of age). (g) The 405 nm control ΔF/F calcium signal (top) and heatmap (bottom) of Esr1^MPOA^ neurons aligned to the onset of the following behaviors: female intruder in, male intruder in, object in, fake pup in and water spray (from left to right). (h-i) The mean AUC of the ΔF/F during various social and non-social behaviors to compare responses across behaviors in hostile virgin males and Trpc5^Esr1^-OE virgin males (h) and responses of the same behavior in hostile virgin males and Trpc5^Esr1^-OE virgin males (i) (n = 4 mice, 12 weeks of age). Data presented as mean ± SEM, p value determined using two-way ANOVA (b,e,h), paired t-test (c), unpaired t-test (f,i).

**Extended data Figure 13.**
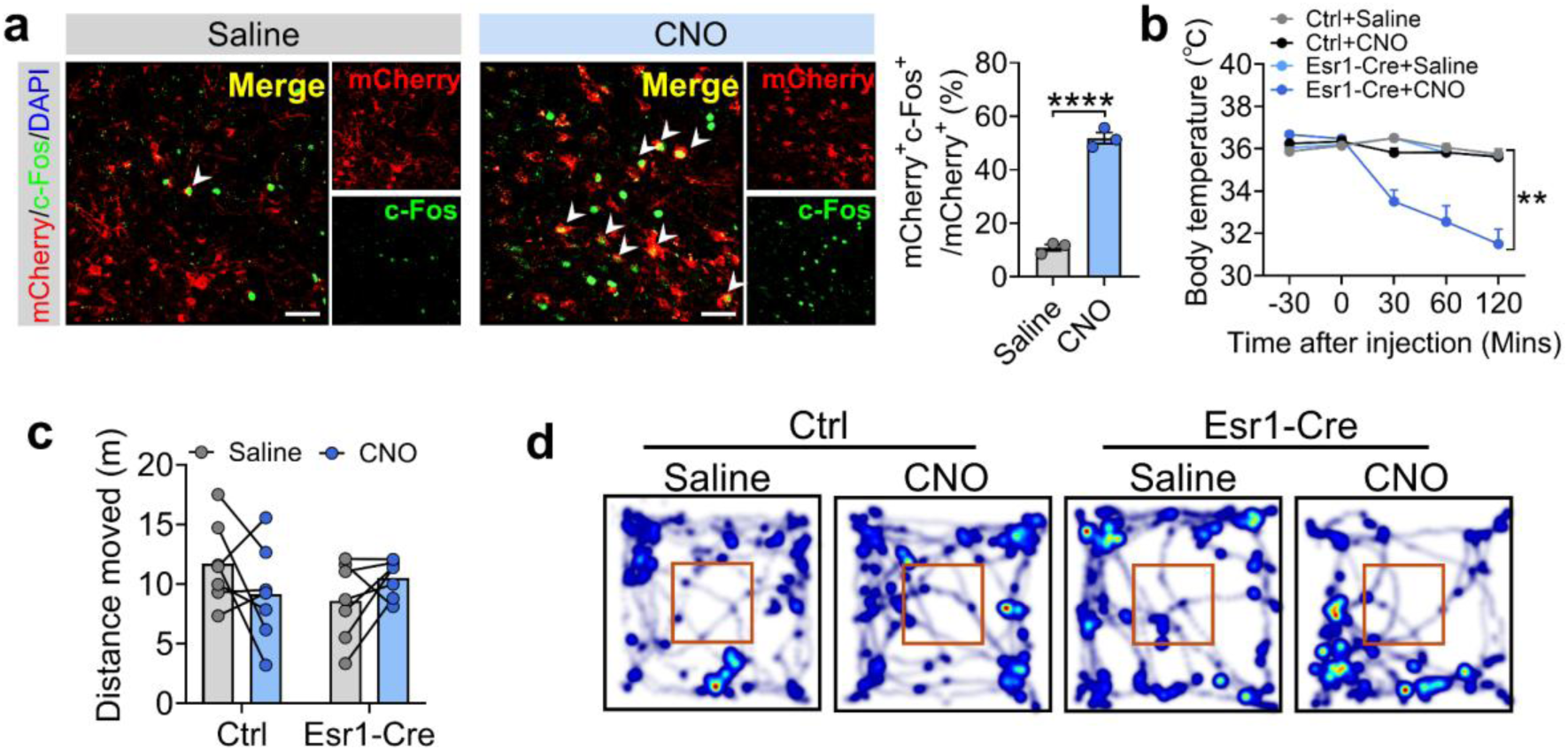
Chemogenetic activation of Esr1^MPOA^ neurons increases pup investigation, reduces body temperature, but does not affect locomotor activity. (a) Left: representative microscopic images showing mCherry (red), c-Fos immunoreactivity (green), and merge in virgin male Esr1-Cre mice infected with AAV-DIO-hM3Dq-mCherry, 30 min after i.p. injections of saline or CNO (0.5 mg kg^-1^). Scale bar: 100 μm. Right: quantification of the percentage of mCherry neurons expressing c-Fos (green) following saline or CNO injection (n = 3 mice per group, 18 weeks of age). (b) The body temperature curve changes in control and Esr1-Cre virgin males treated with saline or CNO (n = 6-7 mice per group, 11 weeks of age). (c-d) The distance moved (c) and heatmap of movement (d) in the open field test between control and Esr1-Cre virgin males treated with saline or CNO (n = 6-7 mice per group, 11 weeks of age). Data presented as mean ± SEM. p value determined using two-way ANOVA (b,c) and unpaired t-test (a). *p < 0.05, **p < 0.01 and ****p < 0.0001.

**Extended data Figure 14.**
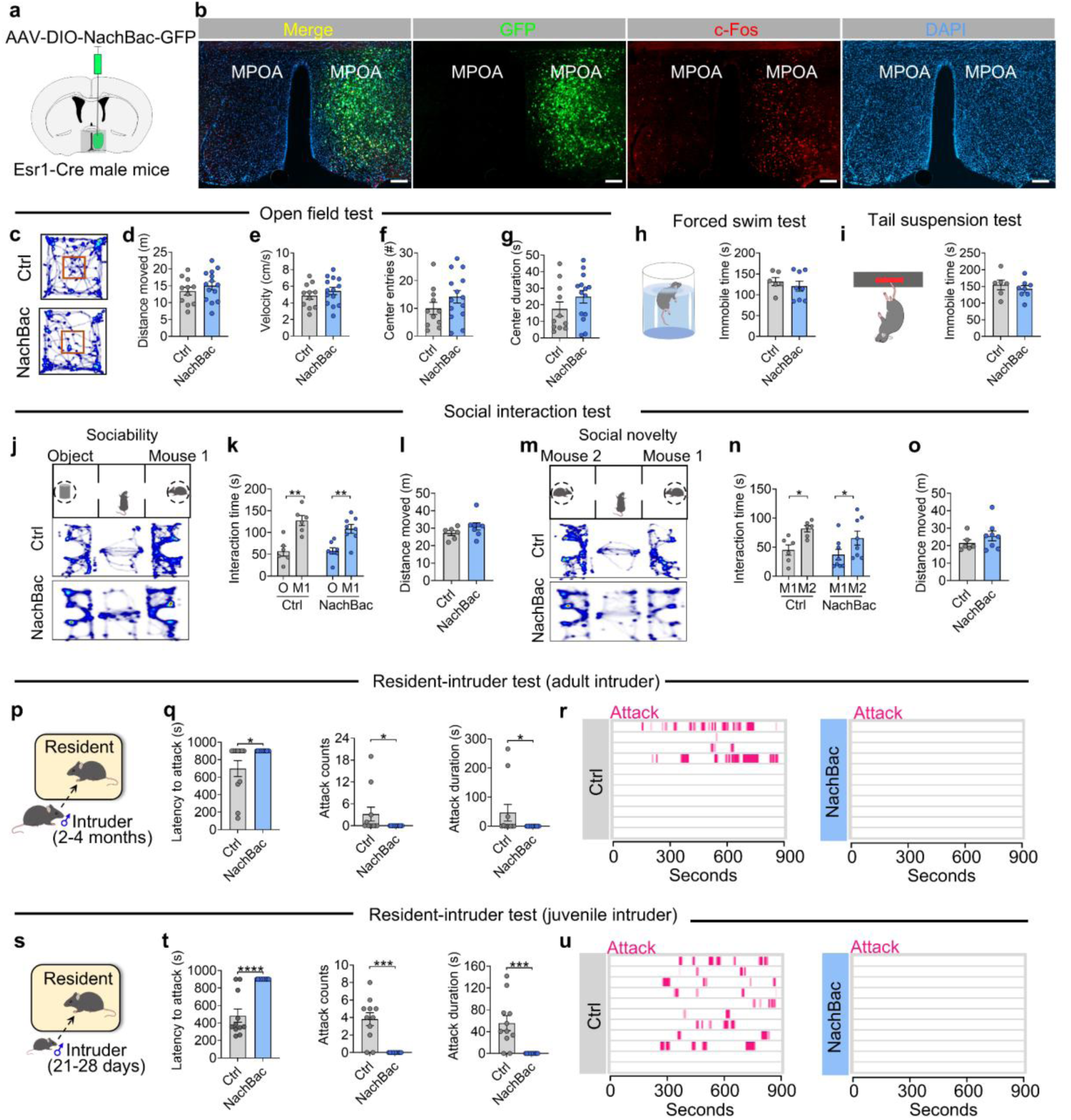
Effects of chronic activation of Esr1^MPOA^ neurons on other social behaviors. (a) Schematic of AAV-DIO-NachBac-GFP injection into the right MPOA of virgin Esr1-Cre male mice. (b) Representative images showing GFP-labeled Esr1^MPOA^ neurons expressing c-Fos in the right side of MPOA in virgin males. The left uninjected side was used as a control. Scale bar, 100 μm. (c-g) Heatmap of movement in the open field test (c), total distance moved (d), velocity (e), center entries (f) and center duration (g) between control and NachBac virgin male mice (n=11 mice per group, 12 weeks of age). (h-i) Immobile time in forced swim test (h) and tail suspension test (i) between control and NachBac virgin male mice (n=6-8 mice per group, 12 weeks of age). (j) Schematic diagram of three-chamber social ability test. (k) Interaction time with object and mouse 1 in chamber between control and NachBac virgin male mice in social ability test (n = 6-8 mice per group, 13 weeks of age). (l) Distances moved in three chambers between control and NachBac virgin male mice in social ability test (n = 6-8 mice per group, 13 weeks of age). (m) Schematic diagram of three-chamber social novelty test. (n) Interaction time with mouse 2 and mouse a in chamber between control and NachBac virgin male mice in social novelty test (n = 6-8 mice per group, 13 weeks of age). (o) Distances moved in three chambers between control and NachBac virgin male mice in social novelty test (n = 6-8 mice per group, 13 weeks of age). (p) Schematic diagram of resident-intruder assay (Group housed adult male intruder mice). (q) Latency to attack (left), number of attacks (middle), and duration of attack (right) between control and NachBac virgin male mice (n = 11 mice per group, 15 weeks of age). (r) Raster plots showing inter-male aggressive behavior. (s) Schematic diagram of resident-intruder assay (Group housed juvenile male intruder mice). (t) Latency to attack (left), number of attacks (middle), and duration of attack (right) between control and NachBac virgin male mice (n = 11 mice per group, 15 weeks of age). (u) Raster plots showing inter-male aggressive behavior. Data presented as mean ± SEM, p value determined using two-way ANOVA (K,N), unpaired t-tests (d,e,f,g,h,i,l,o) and Mann-Whitney test (q,t). *p < 0.05, **p < 0.01, ***p < 0.001 and ****p < 0.0001.

**Extended data Figure 15.**
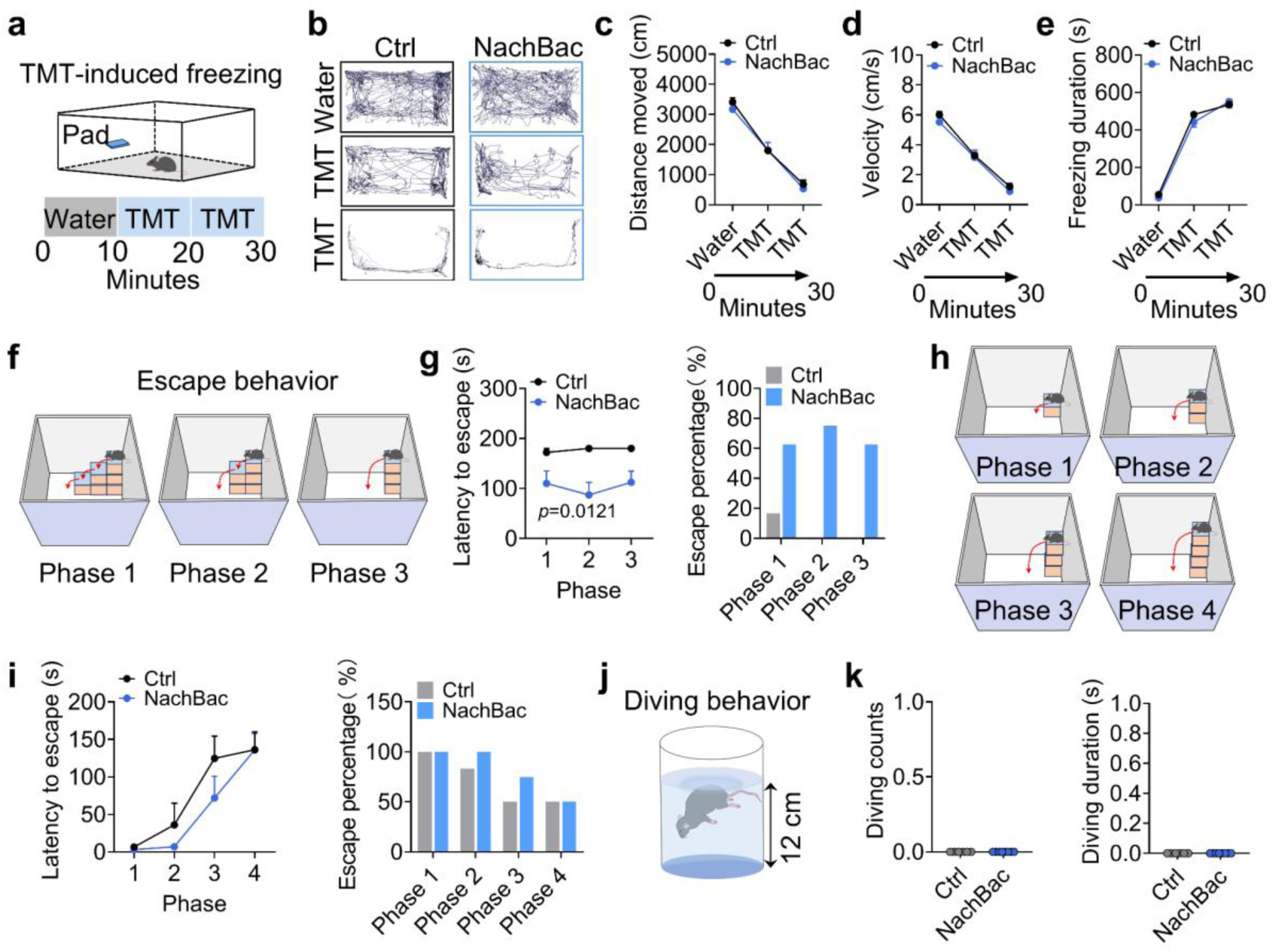
NachBac-induced activation of Esr1^MPOA^ neurons enhances escape behavior but not exploratory diving behavior. (a) Illustration showing TMT-induced behavior in the home cage. Water was first added to the pad for 10 minutes, followed by replacement with a new pad containing TMT for an additional 20 minutes. (b) The track of movement. (c-d) The distance moved (c) and velocity (d) duration within 10 minutes across the water, TMT and TMT exposure (n = 6-8 mice per group, 14 weeks of age). (e) Freezing duration within 10 minutes across the water, TMT, and TMT exposure. (n = 6-8 mice per group; 14 weeks of age). (f) Schematic illustrating a multi-phase escape task with progressively increasing difficulty from phase 1 to 3. (g) Quantification of escape latency (left) and percentage of mice that successfully escaped (right) in control and NachBac virgin male mice (n = 6-8 mice per group; 13 weeks of age). (h) Schematic illustrating a multi-phase escape task with progressively increasing difficulty from phase 1 to 4. (i) Quantification of escape latency (left) and percentage of mice that successfully escaped (right) in control and NachBac virgin male mice (n = 6-8 mice per group; 13 weeks of age). (j) Illustration showing diving behavior in a glass beaker (beaker height: 18 cm; water temperature: 25℃; depth: 12 cm). (k) Diving counts (left) and durations (right) in control and NachBac virgin male mice (n = 6-8 mice per group, 12 weeks of age). Data presented as mean ± SEM, p value determined using two-way ANOVA (c,d,e,j,i) and Mann-Whitney test (k).

## References

1 Feldman, R. The adaptive human parental brain: implications for children’s social development. Trends Neurosci 38, 387–399, doi:10.1016/j.tins.2015.04.004 (2015).

2 Ingram, R. E. & Ritter, J. Vulnerability to depression: cognitive reactivity and parental bonding in high-risk individuals. J Abnorm Psychol 109, 588–596, doi:10.1037//0021-843x.109.4.588 (2000).

3 Frosch, C. A., Schoppe-Sullivan, S. J. & O’Banion, D. D. Parenting and Child Development: A Relational Health Perspective. Am J Lifestyle Med 15, 45–59, doi:10.1177/1559827619849028 (2021).

4 McLeod, J. D. Childhood parental loss and adult depression. J Health Soc Behav 32, 205–220 (1991).

5 Yoo, H. I., Kim, B. N., Shin, M. S., Cho, S. C. & Hong, K. E. Parental attachment and its impact on the development of psychiatric manifestations in school-aged children. Psychopathology 39, 165–174, doi:10.1159/000092677 (2006).

6 Stockley, P. & Hobson, L. Paternal care and litter size coevolution in mammals. Proc Biol Sci 283, doi:10.1098/rspb.2016.0140 (2016).

7 Rogers, F. D. & Bales, K. L. Mothers, Fathers, and Others: Neural Substrates of Parental Care. Trends Neurosci 42, 552–562, doi:10.1016/j.tins.2019.05.008 (2019).

8 Dulac, C., O’Connell, L. A. & Wu, Z. Neural control of maternal and paternal behaviors. Science 345, 765–770, doi:10.1126/science.1253291 (2014).

9 Rilling, J. K. & Young, L. J. The biology of mammalian parenting and its effect on offspring social development. Science 345, 771–776, doi: 10.1126/science.1252723 (2014).

10 Kohl, J. & Dulac, C. Neural control of parental behaviors. Curr Opin Neurobiol 49, 116–122, doi:10.1016/j.conb.2018.02.002 (2018).

11 Inada, K. et al. Plasticity of neural connections underlying oxytocin-mediated parental behaviors of male mice. Neuron 110, 2009–2023 e2005, doi:10.1016/j.neuron.2022.03.033 (2022).

12 Lee, A. W. & Brown, R. E. Medial preoptic lesions disrupt parental behavior in both male and female California mice (Peromyscus californicus). Behav Neurosci 116, 968–975, doi:10.1037//0735-7044.116.6.968 (2002).

13 Kohl, J. et al. Functional circuit architecture underlying parental behaviour. Nature 556, 326–331, doi:10.1038/s41586-018-0027-0 (2018).

14 Wu, Z., Autry, A. E., Bergan, J. F., Watabe-Uchida, M. & Dulac, C. G. Galanin neurons in the medial preoptic area govern parental behaviour. Nature 509, 325–330, doi:10.1038/nature13307 (2014).

15 Yoshihara, C. et al. Calcitonin receptor signaling in the medial preoptic area enables risk-taking maternal care. Cell Rep 35, 109204, doi:10.1016/j.celrep.2021.109204 (2021).

16 Alcantara, I. C. et al. A hypothalamic circuit that modulates feeding and parenting behaviours. Nature, doi:10.1038/s41586-025-09268-5 (2025).

17 Stagkourakis, S. et al. A Neuro-hormonal Circuit for Paternal Behavior Controlled by a Hypothalamic Network Oscillation. Cell 182, 960–975 e915, doi:10.1016/j.cell.2020.07.007 (2020).

18 Zhang, G. W. et al. Medial preoptic area antagonistically mediates stress-induced anxiety and parental behavior. Nat Neurosci 24, 516–528, doi:10.1038/s41593-020-00784-3 (2021).

19 Fang, Y. Y., Yamaguchi, T., Song, S. C., Tritsch, N. X. & Lin, D. A Hypothalamic Midbrain Pathway Essential for Driving Maternal Behaviors. Neuron 98, 192–207 e110, doi:10.1016/j.neuron.2018.02.019 (2018).

20 Rosenblatt, J. S. & Ceus, K. Estrogen implants in the medial preoptic area stimulate maternal behavior in male rats. Horm Behav 33, 23–30, doi:10.1006/hbeh.1997.1430 (1998).

21 Smiley, K. O., Brown, R. S. E. & Grattan, D. R. Prolactin Action Is Necessary for Parental Behavior in Male Mice. J Neurosci 42, 8308–8327, doi:10.1523/JNEUROSCI.0558-22.2022 (2022).

22 Clapham, D. E. TRP channels as cellular sensors. Nature 426, 517–524, doi:10.1038/nature02196 (2003).

23 Li, Y. et al. Loss of transient receptor potential channel 5 causes obesity and postpartum depression. Cell 187, 4176–4192 e4117, doi: 10.1016/j.cell.2024.06.001 (2024).

24 Isogai, Y. et al. Multisensory Logic of Infant-Directed Aggression by Males. Cell 175, 1827–1841 e1817, doi:10.1016/j.cell.2018.11.032 (2018).

25 Ammari, R. et al. Hormone-mediated neural remodeling orchestrates parenting onset during pregnancy. Science 382, 76–81, doi:10.1126/science.adi0576 (2023).

26 Wang, W. et al. Coordination of escape and spatial navigation circuits orchestrates versatile flight from threats. Neuron 109, 1848–1860 e1848, doi:10.1016/j.neuron.2021.03.033 (2021).

27 Corona, A., Choe, J., Munoz-Castaneda, R., Osten, P. & Shea, S. D. A circuit from the locus coeruleus to the anterior cingulate cortex modulates offspring interactions in mice. Cell Rep 42, 112771, doi:10.1016/j.celrep.2023.112771 (2023).

28 Yoshihara, C., Numan, M. & Kuroda, K. O. Oxytocin and Parental Behaviors. Curr Top Behav Neurosci 35, 119–153, doi:10.1007/7854_2017_11 (2018).

29 Parent, C. I., Del Corpo, A., Cameron, N. M. & Meaney, M. J. Maternal care associates with play dominance rank among adult female rats. Dev Psychobiol 55, 745–756, doi:10.1002/dev.21070 (2013).

30 Bridges, R. S. Neuroendocrine regulation of maternal behavior. Front Neuroendocrinol 36, 178–196, doi:10.1016/j.yfrne.2014.11.007 (2015).

31 Cirulli, F., Berry, A. & Alleva, E. Early disruption of the mother-infant relationship: effects on brain plasticity and implications for psychopathology. Neurosci Biobehav Rev 27, 73–82, doi:10.1016/s0149-7634(03)00010-1 (2003).

32 Hofferth, S. & Lee, Y. Family structure and trends in US fathers’ time with children, 2003-2013. Fam Sci 6, 318–329, doi:10.1080/19424620.2015.1082805 (2015).

33 Wray, D., Ingenfeld, J., Milkie, M. A. & Boeckmann, I. Beyond childcare: Changes in the amount and types of parent-child time over three decades. Can Rev Sociol 58, 327–351, doi:10.1111/cars.12356 (2021).

34 Jackson, D. B., Newsome, J. & Beaver, K. M. Does early paternal involvement predict offspring developmental diagnoses? Early Hum Dev 103, 9–16, doi:10.1016/j.earlhumdev.2016.07.001 (2016).

35 Viellard, J. M. A. et al. A subiculum-hypothalamic pathway functions in dynamic threat detection and memory updating. Curr Biol 34, 2657–2671 e2657, doi:10.1016/j.cub.2024.05.006 (2024).

36 Harikai, N., Sugawara, T., Tomogane, K., Mizuno, K. & Tashiro, S. Acute heat stress induces jumping escape behavior in mice. Physiol Behav 83, 373–376, doi:10.1016/j.physbeh.2004.06.019 (2004).

37 Barik, A., Thompson, J. H., Seltzer, M., Ghitani, N. & Chesler, A. T. A Brainstem-Spinal Circuit Controlling Nocifensive Behavior. Neuron 100, 1491–1503 e1493, doi:10.1016/j.neuron.2018.10.037 (2018).

38 Zholos, A. V. Trpc5. Handb Exp Pharmacol 222, 129–156, doi:10.1007/978-3-642-54215-2_6 (2014).

39 Broker-Lai, J. et al. Heteromeric channels formed by TRPC1, TRPC4 and TRPC5 define hippocampal synaptic transmission and working memory. EMBO J 36, 2770–2789, doi:10.15252/embj.201696369 (2017).

40 Ordaz, B. et al. Calmodulin and calcium interplay in the modulation of TRPC5 channel activity. Identification of a novel C-terminal domain for calcium/calmodulin-mediated facilitation. J Biol Chem 280, 30788–30796, doi:10.1074/jbc.M504745200 (2005).

41 Kinoshita-Kawada, M. et al. Inhibition of TRPC5 channels by Ca2+-binding protein 1 in Xenopus oocytes. Pffugers Arch 450, 345–354, doi:10.1007/s00424-005-1419-1 (2005).

42 Ray, S. et al. An Examination of Dynamic Gene Expression Changes in the Mouse Brain During Pregnancy and the Postpartum Period. G3 (Bethesda) 6, 221–233, doi:10.1534/g3.115.020982 (2015).

43 Bukhari, S. A. et al. Neurogenomic insights into paternal care and its relation to territorial aggression. Nat Commun 10, 4437, doi:10.1038/s41467-019-12212-7 (2019).

44 Seelke, A. M. H. et al. Fatherhood alters gene expression within the MPOA. Environ Epigenet 4, dvy026, doi:10.1093/eep/dvy026 (2018).

45 Qu, N. et al. A POMC-originated circuit regulates stress-induced hypophagia, depression, and anhedonia. Mol Psychiatry 25, 1006–1021, doi:10.1038/s41380-019-0506-1 (2020).

